# POLO kinase inhibits Protein Phosphatase 1 to promote the Spindle Assembly Checkpoint and prevent aneuploidy

**DOI:** 10.1101/2024.12.23.630043

**Authors:** Margarida Moura, João Barbosa, Inês Pinto, Nelson Leça, Sofia Cunha-Silva, Arianna Esposito Verza, Paulo D. Pedroso, Sarah Lemaire, Joana Duro, Miguel Silva, André Oliveira, Ricardo Reis, Jakob Nilsson, Claudio Sunkel, Andrea Musacchio, Mathieu Bollen, Carlos Conde

## Abstract

Protein phosphatase 1 (PP1) is essential for spindle assembly checkpoint (SAC) silencing and mitotic exit, but its regulation during mitosis remains ill-defined. Here, we demonstrate *in vitro* and in *Drosophila* cells that the mitotic kinase POLO phosphorylates PP1α87B at a conserved residue (T286) within a pocket implicated in the recognition of RVxF-containing target proteins. Phosphorylation of T286 inhibits PP1α87B binding to the RVxF motif of the SAC kinase MPS1, dampening the dephosphorylation of MPS1 T-loop. Phosphorylation of T286 is dynamically regulated during mitosis. It occurs at unattached/tensionless kinetochores and decreases as chromosomes congress. Expression of phosphomimetic PP1α87B^T286D^ prevents MPS1 inactivation in metaphase and causes a SAC-dependent delay of anaphase onset. Conversely, an unphosphorylatable PP1α87B^T286A^ mutant impairs MPS1 activation at unattached kinetochores and weakens the SAC. *In vivo*, larval neuroblasts expressing PP1α87B^T286^ phosphomutants exhibit frequent chromosome mis-segregation and aneuploidy. Thus, our findings identify POLO-mediated phosphorylation of PP1α87B as a critical regulatory strategy that fine-tunes phosphatase activity to ensure a robust and timely SAC and prevent genome instability.

## INTRODUCTION

To ensure accurate genome partitioning, dividing eukaryotic cells halt anaphase onset until all kinetochores are bound to spindle microtubules (Lara-Gonzalez et al., 2021; McAinsh and Kops, 2023). This surveillance mechanism is known as the Spindle Assembly Checkpoint (SAC) and its downstream target is the Anaphase Promoting Complex/Cyclosome (APC/C), an E3 ubiquitin ligase that targets SECURIN and CYCLIN B for proteasome-mediated proteolysis. Degradation of SECURIN and CYCLIN B enables sister chromatid separation and CYCLIN-DEPENDENT KINASE 1 (CDK1) inactivation, events required for chromosome segregation and mitotic exit (Lara-Gonzalez et al., 2021; McAinsh and Kops, 2023). Both a weak and an overactive SAC can lead to chromosome mis-segregation, culminating in aneuploidy, a condition frequently associated with tumorigenesis (Michel et al., 2001; Iwanaga et al., 2007; Morais da Silva et al., 2013; Resende et al., 2018). To safeguard mitotic fidelity, the SAC evolved to be simultaneously robust and responsive. These SAC features ensure that mitotic exit is delayed when even a single kinetochore remains unattached to microtubules, while enabling prompt anaphase onset as soon as all kinetochore attachments are established (Corno et al., 2023). A highly dynamic tug-of-war between kinases and phosphatases at kinetochores regulates the SAC (Saurin, 2018; Moura and Conde, 2019). Timely SAC silencing relies largely on Protein Phosphatase 1 (PP1), a conserved serine/threonine phosphatase that dephosphorylates the MELT motifs of KNL1 at metaphase kinetochores (London et al., 2012; Zhang et al., 2014; Nijenhuis et al., 2014) and the T-loop of MPS1, causing inactivation of this crucial SAC kinase (Moura et al., 2017). Together, these events limit the accumulation of SAC proteins at kinetochores, switch off the SAC, and enable swift onset of anaphase. However, it remains unclear how PP1 is precisely controlled to prevent premature checkpoint silencing while ensuring sufficient activity for timely anaphase onset upon chromosome biorientation. AURORA B-mediated phosphorylation of the PP1-binding SILK and RVxF-type short linear motifs (SLiMs) of KNL1 has been shown to hinder the recruitment of PP1 to unattached or tensionless kinetochores (Liu et al., 2010; Rosenberg et al., 2011; Bajaj et al., 2018; Nasa et al., 2018). However, rendering these SLiMs unphosphorylatable, by mutation to AILK and RVAF, only marginally reduced (by 25%) MELT phosphorylation in nocodazole-treated cells (Nijenhuis et al., 2014), indicating that additional mechanisms limit PP1 activity at kinetochores during early mitosis. Additional layers for controlling PP1 activity may involve the formation of specific holoenzymes through association of PP1 with Regulatory Interactors of Protein Phosphatase One (RIPPOs) (Bollen et al. 2010; Cao et al., 2022) and the phosphorylation of a conserved threonine residue in the disordered C-terminal tail of PP1. This threonine (T320 in PP1α, T316 in PP1β and T311 in PP1γ) is a substrate for CDK2 and CDK1, and has been associated with an inhibitory effect on PP1 during the G1/S transition and mitosis, respectively (Dohadwala et al., 1994; Yamano et al., 1994; Kwon et al., 1997; Liu et al., 1999; Wu et al., 2009; Grallert et al., 2015; Qian et al., 2015). Structural, semisynthetic and biochemical data indicate that T320/316/311 phosphorylation inhibits PP1 function by interfering with the binding to RIPPOs rather than by direct structural changes or substrate competition (Salvi et al., 2020). However, the mechanism by which this phosphorylation controls holoenzyme assembly to reduce PP1 activity and its relevance for SAC signaling remains unclear. SAC silencing precedes APC/C-mediated ubiquitination of CYCLIN B, which implies that PP1-dependent dephosphorylation of KNL1 and MPS1 occurs when CDK1 is still active. Furthermore, PP1 also triggers prompt APC/C-dependent degradation of CYCLIN B downstream of the SAC through dephosphorylation of the N-terminus of CDC20 (Kim et al., 2017; Bancroft et al., 2020). This event contributes to accelerating the metaphase to anaphase transition and, as for the dephosphorylation of KNL1 and MPS1, takes place when PP1 is still largely phosphorylated at T320/T316/T311 (Bancroft et al., 2020). These observations challenge the involvement of CDK1:CYCLIN B-mediated inhibition of PP1 in controlling timely anaphase onset.

Here, we set out to investigate the regulation of PP1 in mitosis by using *Drosophila melanogaster* as a model organism. The genome of fruit flies has four loci encoding PP1 isoforms, of which PP1α87B is the most abundant and responsible for the bulk of PP1 activity during mitosis (Axton et al., 1990; Dombrádi et al. 1990; Moura et al., 2017; Audett et al., 2022). Importantly, PP1α87B has no C-terminal tail, and hence, lacks the regulatory CDK-targeted threonine present in most PP1 isoforms. We took advantage of this naturally occurring simpler system to explore hitherto uncovered mechanisms that regulate PP1 in mitosis.

## RESULTS

### AURORA B- and CDK1-mediated mechanisms do not fully account for the inhibition of PP1α87B’s SAC-antagonizing activity

We first examined to which extent previously described PP1-inhibitory mechanisms also apply to *Drosophila* S2 cells. AURORA B phosphorylates residues within or flanking the RVxF and SILK motifs of KNL1 at unattached/tensionless kinetochores, which decreases their binding affinity for PP1. This limits PP1 recruitment to the outer kinetochore before stable attachments to spindle microtubules are established (Liu et al., 2010; Rosenberg et al., 2011; Bajaj et al., 2018; Nasa et al., 2018). This mechanism also operates in S2 cells: AURORA B phosphorylates the RVxF motif of SPC105R (*Drosophila* KNL1 orthologue) in the absence of intra-kinetochore tension (Drpic et al., 2015; Barbosa et al., 2020) and chemical inhibition of AURORA B enhances the accumulation of PP1α87B at unattached kinetochores (Moura et al., 2017). To test whether this AURORA B-controlled recruitment of PP1α87B is sufficient to prevent premature SAC silencing, we constitutively tethered PP1α87B to the outer kinetochore of S2 cells by fusing it to the kinetochore protein MIS12 and monitored mitotic progression by live cell imaging. Compared to MIS12-EGFP, expression of the MIS12-EGFP-PP1α87B transgene did not accelerate anaphase onset in asynchronous cultured cells or in cells treated with colchicine to induced microtubule depolymerization (Figure 1A). Importantly, expression of MIS12-EGFP-PP1α87B restored timely mitotic exit in S2 cells depleted of endogenous PP1α87B, demonstrating that the kinetochore-tethered phosphatase was functional (Figure 1A). These results show that bypassing the phospho-regulation of PP1-docking motifs on SPC105R is not sufficient to silence or weaken the SAC.

**Figure 1.**
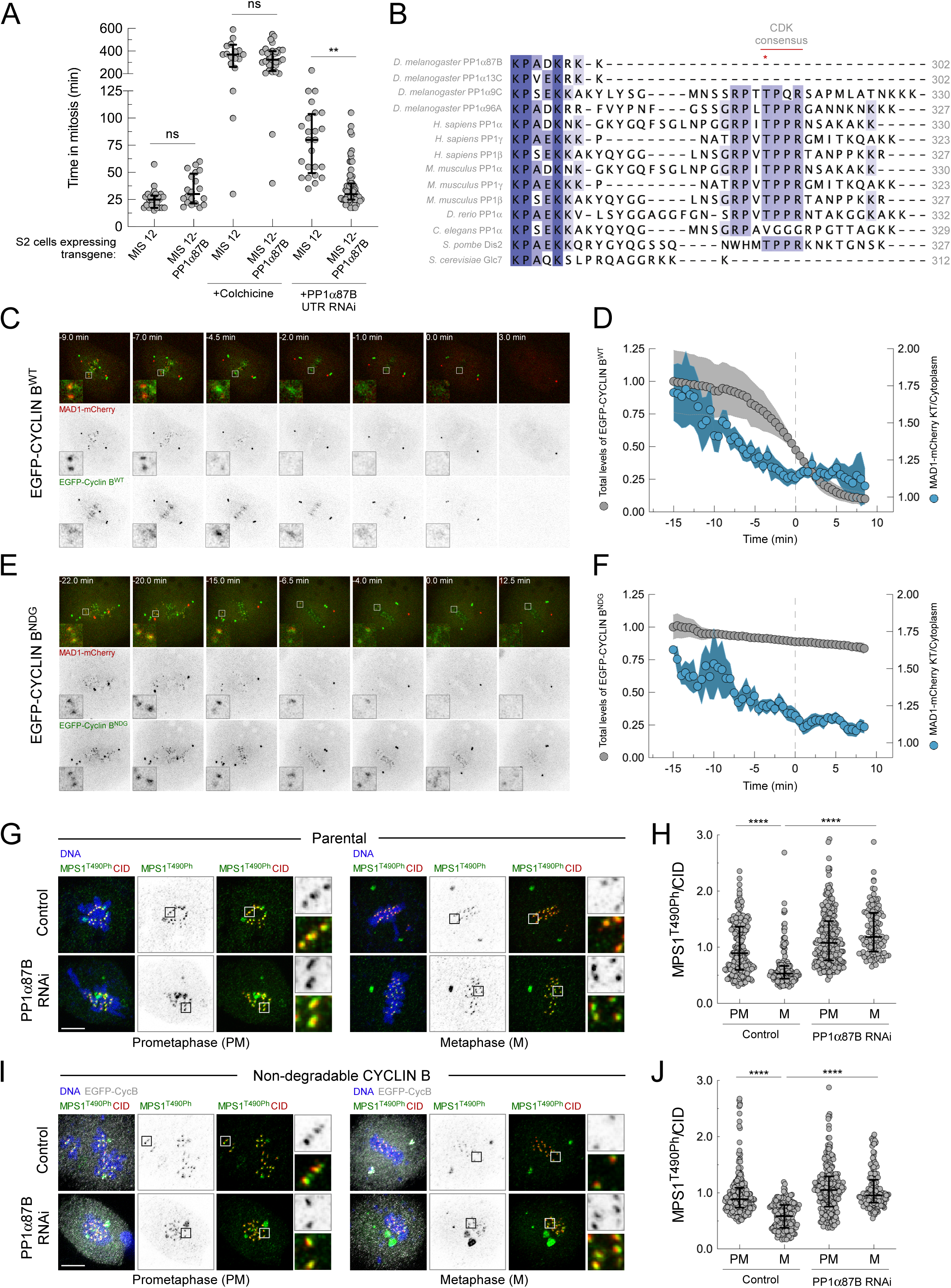
PP1α87B is impervious to CDK1 activity and fails to antagonize the SAC when forced to accumulate at unattached or prometaphase kinetochores. (A) Mitotic duration of *Drosophila* S2 cells expressing the indicated transgenes in the indicated conditions. Mitotic progression was monitored by time-lapse microscopy. The mitotic duration was defined by the length of time measured between nuclear envelope breakdown and anaphase onset or chromatin decondensation in the case of colchicine-treated cells (n ≥ 20 cells for each condition). (B) Clustal W alignments of the C-terminus protein sequence of indicated PP1 orthologues. (C,D) Mitotic progression of S2 cells expressing MAD1-mCherry and EGFP-Cyclin B^WT^ monitored by time-lapse microscopy (C) and quantification of MAD1-mCherry kinetochore levels and EGFP-Cyclin B^WT^ total cellular levels through time (D). Time 0 corresponds to anaphase onset (n = 8 cells). (E,F) Mitotic progression of S2 cells expressing MAD1-mCherry and EGFP-Cyclin B^NDG^ monitored by time-lapse microscopy (E) and quantification of MAD1-mCherry kinetochore levels and EGFP-Cyclin B^NDG^ total cellular levels through time (F). Because these cells often fail to undergo anaphase, Time 0 corresponds to the first frame where maximal inter-kinetochore distance was achieved, as determined based on EGFP-Cyclin B^NDG^ signal for kinetochore reference (n = 10 cells). (G-J) Representative immunofluorescence images (G,I) and corresponding quantifications of MPS1 T490 phosphorylation (MPS1^T490Ph^) levels at kinetochores (H,J) of control and PP1α87B**-**depleted S2 cells in prometaphase and metaphase (n ≥ 112 kinetochores for each condition). Parental S2 cells (G,H) or cells expressing EGFP-Cyclin B^NDG^ (I,J) were analyzed for MPS1^T490Ph^ fluorescence intensity relative to CID signal. Data information: in A, H and J, data are presented as median with interquartile range; asterisks indicate that differences between mean ranks are statistically significant, **p<0.01; ****p<0.0001; ns, non-significant (Kruskal-Wallis, Dunn’s multiple comparison test). Scale bars: 5μm. See also Figure S1.

Phosphorylation of PP1 C-terminal tail by CDK1 has been proposed to negatively regulate PP1 activity in different systems (Wu et al., 2009; Grallert et al., 2015; Qian et al., 2015). However, PP1α87B has no C-terminal tail and therefore lacks the regulatory CDK-targeted threonine (Figure 1B). In contrast, the *Drosophila* paralogues PP1α9C and PP1α96A do have a disordered C-terminal tail with a threonine in a consensus motif (PxTPP) for phosphorylation by CDK1 (Figure 1B). This prompted us to test whether CDK1 activity contributes to reducing overall PP1 function in mitotic cells. We generated S2 cell lines that stably co-express MAD1-mCherry and EGFP-tagged wild type (WT) or non-degradable (NDG) CYCLIN B (Sigrist et al., 1995), and monitored the kinetochore localization of MAD1-mCherry as a readout of SAC signaling (Figure 1C-F). In EGFP-CYCLIN B^WT^-expressing cells, the MAD1-mCherry signal at kinetochore pairs began to fade as these aligned to the metaphase plate (∼5 min before anaphase onset) and equaled background intensity ∼2 min prior to the onset of anaphase, when ∼75% of EGFP-CYCLIN B^WT^ levels were still present in the cell (Figure 1C,D and Video 1). This indicates that the SAC is switched off (MAD1-mCherry absent from kinetochores) even under elevated CDK1:CYCLIN B activity (indirectly inferred from EGFP-CYCLIN B^WT^ levels). This notion was further supported by our finding that the kinetics of MAD1-mCherry clearance from bioriented kinetochores was not affected by deficient CYCLIN B proteolysis in cells expressing EGFP-CYCLIN B^NDG^ (Figure 1E,F and Video 2).

To examine whether SAC silencing in the presence of high CDK1:CYCLIN B levels was PP1α87B-dependent, we used a phospho-specific antibody (Jelluma et al., 2008) to assess the phosphorylation of MPS1 T-loop (T490Ph) at kinetochores. MPS1^T490Ph^ is an established PP1α87B substrate whose dephosphorylation during metaphase is required for MPS1 inactivation and SAC silencing in *Drosophila* cells (Moura et al., 2017). Immunofluorescence analysis showed that the expression of EGFP-CYCLIN B^NDG^ did not prevent the inactivation of MPS1 in metaphase, as revealed by the substantial decrease in MPS1 T-loop phosphorylation as cells progressed from prometaphase to metaphase (Figure 1G-J). The dephosphorylation of MPS1 at T490 was also unaffected in metaphase cells treated with MG132, which inhibits proteasome-mediated degradation of CYCLIN A and CYCLIN B (Figure S1A,B). Hence, the inactivation of MPS1 in MG132-treated and EGFP-CYCLIN B^NDG^-expressing cells mirrors what is observed in untreated control parental S2 cells and is mediated by PP1α87B, as its depletion impaired the dephosphorylation of MPS1^T490Ph^ at metaphase kinetochores (Figure 1G-J and Figure S1A,B). In accordance with the activation status of MPS1, the phosphorylation of MAD1 C-terminus at T726 (T716 in human MAD1), which is catalyzed by MPS1 and rate-limiting for SAC signaling (Faesen et al., 2017; Ji et al., 2017; Ji et al., 2018; Allan et al., 2020; Fischer et al., 2022), also decreased at metaphase kinetochores of cells impaired in CYCLIN A and CYCLIN B degradation unless PP1α87B was depleted (Figure S1C-K). Collectively, these results indicate that inactivation of MPS1 by PP1α87B can still occur when high levels of CYCLIN A and CYCLIN B are present in the cell, showing that CDK1 activity, at best, contributes marginally to prevent PP1α87B from silencing the SAC in *Drosophila* S2 cells.

### POLO phosphorylates PP1α87B at unattached/tensionless kinetochores

Given that PP1α87B is unlikely to be phosphorylated by CDK1 (lacks CDK1-regulated C-terminal tail) and its forced accumulation at kinetochores (fusion to MIS12) fails to prematurely silence the SAC (Figure 1A), we hypothesized that other mechanisms operate during prometaphase to restrain PP1α87B activity. The presence of a conserved consensus sequence for a POLO/PLK1 phosphorylation site (T286) in the RVxF-binding pocket of PP1α87B (Figure 2A and 2B) prompted us to interrogate whether POLO/PLK1 could directly regulate PP1α87B/PP1 activity in mitosis. First, we assessed PLK1 capacity to phosphorylate PP1. We raised an antibody against phosphorylated T286 (T288 in human PP1γ) and performed *in vitro* kinase assays with recombinant purified proteins. Immunoblotting showed an increase of 6xHis-PP1γ^WT^ phosphorylation upon incubation with 6xHis-PLK1, which was impaired after T288 was converted to alanine (6xHis-PP1γ^T288A^), indicating that PLK1 can phosphorylate PP1γ at T288 *in vitro* (Figure 2C).

**Figure 2.**
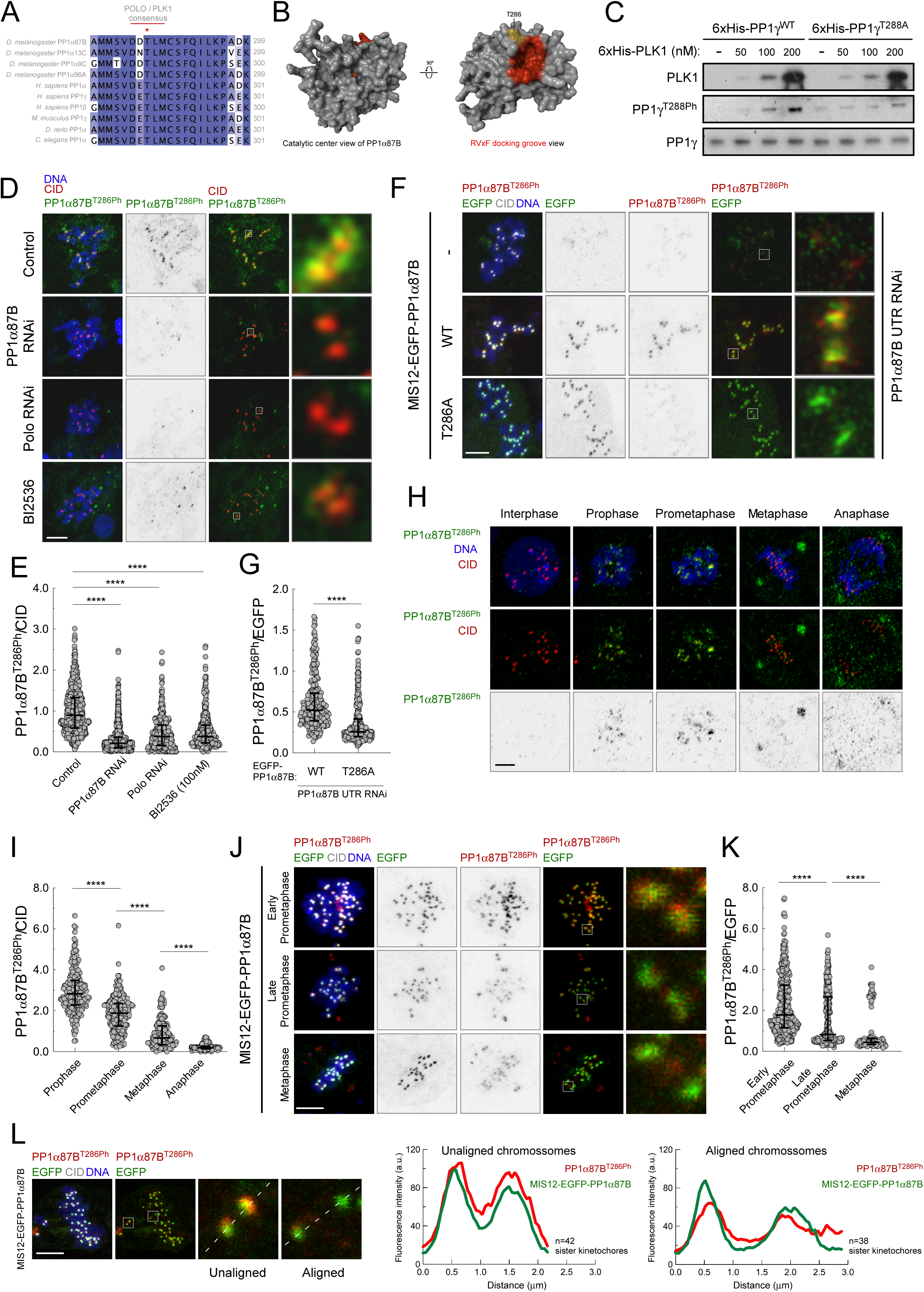
POLO phosphorylates a conserved threonine in the RVxF-binding pocket of PP1α87B at kinetochores of unaligned chromosomes. (A) Clustal W alignments of the C-terminus protein sequence of indicated PP1 orthologues. The conserved POLO/PLK1 consensus motif and putative phosphorylatable threonine are indicated by the red line and *, respectively. (B) Schematic representation of *Drosophila* PP1α87B (P12982) structure predicted by AlphaFold3. The RVxF docking groove is highlighted in red and the putative POLO phosphorylation site T286 is highlighted in yellow. Fe^3+^ (orange) and the Zn^2+^ (light purple) metals in the catalytic center are represented. (C) *In vitro* kinase assay with increasing concentrations of recombinant 6xHis-PLK1 and 6xHis-PP1γ (450 nM) in the presence of ATP for 30 min. Phosphorylation of T288 was detected by western blot using a phospho-specific antibody raised against the phosphorylated residue. Wild type (WT) and unphosphorylatable (T288A) versions of PP1γ were used as substrates. (D,E) Representative immunofluorescence images (D) and corresponding quantifications (E) of PP1α87B T286 phosphorylation (PP1α87B^T286Ph^) at unattached kinetochores of colchicine-treated S2 cells in the indicated conditions. PP1α87B^T286Ph^ fluorescence intensity was determined relative to CID signal (n ≥ 337 kinetochores for each condition). (F,G) Representative immunofluorescence images (F) and corresponding quantifications (G) of PP1α87B T286 phosphorylation (PP1α87B^T286Ph^) at unattached kinetochores of colchicine-treated S2 cells in the indicated conditions. PP1α87B^T286Ph^ fluorescence intensity was determined relative to EGFP signal (n ≥ 286 kinetochores for each condition). (H,I) Representative immunofluorescence images (H) and corresponding quantifications (I) of PP1α87B T286 phosphorylation (PP1α87B^T286Ph^) levels at kinetochores of S2 cells throughout mitosis. PP1α87B^T286h^ fluorescence intensity was determined relative to CID signal (n ≥ 150 kinetochores for each condition). (J,K) Representative immunofluorescence images (J) and corresponding quantifications (K) of PP1α87B T286 phosphorylation (PP1α87B^T286Ph^) levels at kinetochores of S2 cells expressing Mis12-EGFP-PP1α87B^WT^ in the indicated mitotic stages. PP1α87B^T286h^ fluorescence intensity was determined relative to EGFP signal (n 112 ≥ kinetochores for each condition). (L) Representative image and corresponding quantifications of PP1α87B^T286h^ levels at kinetochore pairs of unaligned and aligned chromosomes of S2 cells expressing Mis12-EGFP-PP1α87B^WT^. Graphs in L represent the mean fluorescence intensity profiles of PP1α87B^T286h^ (red) and Mis12-EGFP-PP1α87B^WT^ (green) along sister-kinetochores (n ≥ 38 sister-kinetochores for each condition). Data information: in E, G, I and K, data are presented as median with interquartile range; asterisks indicate that differences between mean ranks are statistically significant, ****p<0.0001; (Kruskal-Wallis, Dunn’s multiple comparison test in E, I and K and Mann-Whitney test in G). Scale bars: 5μm. See also Figure S2.

To test if POLO phosphorylates PP1α87B at T286 in *Drosophila* cells, we performed immunofluorescence analysis using the phospho-specific antibody. Phosphorylation of PP1α87B at T286 (T286Ph) was uniformly detected at kinetochores of colchicine-treated cells (Figure 2D,E). However, kinetochore levels of T286Ph were severely reduced following RNAi-mediated depletion of POLO or by chemical inhibition of its kinase activity with BI2536 (Figure 2D,E). Depleting PP1α87B or replacing the endogenous protein with a kinetochore-tethered unphosphorylatable mutant for T286 (MIS12-EGFP-PP1α87B^T286A^) also compromised T286Ph staining, which further validated the specificity of the antibody in S2 cells (Figure 2D-G). Collectively, these results strongly support that POLO phosphorylates PP1α87B at T286 in a cellular context.

Next, we assessed how the phosphorylation levels of PP1α87B at T286 evolved during mitosis in asynchronous cultured S2 cells. The kinetochore signal of PP1α87B^T286Ph^ was readily detected in prophase and prometaphase cells but became significantly reduced in metaphase and virtually undetectable in cells undergoing anaphase (Figure 2H,I). Because kinetochore recruitment of PP1α87B increases as chromosome biorientation is established (Moura et al., 2017; Audett et al., 2022), the decrease in T286 phosphorylation at metaphase kinetochores cannot be explained by reduced kinetochore levels of PP1α87B. To validate this conclusion, we assessed the phosphorylation of T286 in S2 cells expressing the outer kinetochore-tethered version of PP1α87B. Kinetochore levels of MIS12-EGFP-PP1α87B remain constant throughout mitosis, thus allowing us to monitor T286Ph relative to the amount of PP1α87B. Immunofluorescence analysis showed that the phosphorylation of T286 was elevated at tensionless kinetochores of unaligned chromosomes, but was strikingly reduced at kinetochore pairs of chromosomes that had already congressed to the metaphase plate and exhibited inter-kinetochore tension (Figure 2J-L). This phosphorylation pattern correlates with the activation status of POLO at kinetochores throughout mitosis (Conde et al., 2013; Barbosa et al., 2020). Notably, in contrast to S2 cells expressing endogenous POLO or a wild type POLO transgene (POLO^WT^-EGFP), preventing POLO inactivation through the expression of a constitutively active version of the kinase (POLO^T182D^-EGFP) was sufficient to retain elevated levels of PP1α87B^T286Ph^ at metaphase kinetochores (Figure S2A,B). Collectively, these data demonstrate that POLO phosphorylates the conserved T286 of PP1α87B and this preferentially occurs at kinetochores of unattached and unaligned chromosomes.

### Phosphorylation of PP1α87B at T286 is required for robust SAC signaling

To dissect the relevance of POLO-mediated phosphorylation of PP1α87B at T286, we generated S2 cell lines expressing EGFP-tagged wild type (WT), unphosphorylatable (T286A) and phosphomimetic (T286D) versions of PP1α87B. Given the higher levels of T286 phosphorylation at unattached/tensionless kinetochores, we first evaluated its importance for SAC signaling. For that purpose, we assessed by live-cell imaging the ability of PP1α87B phosphomutants to arrest in mitosis when incubated with colchicine (Figure 3A). In line with a proficient SAC, cells expressing EGFP-PP1α87B^WT^ or EGFP-PP1α87B^T286D^ significantly delayed the transition to anaphase in the presence of colchicine (∼462 min and ∼470 min, respectively). In contrast, EGFP-PP1α87B^T286A^ cells failed to sustain this arrest as efficiently, exiting mitosis after ∼350 min (Figure 3A). This weakened SAC correlated with a reduction in MPS1 activation (Figure 3B,C) and MPS1-mediated phosphorylation of MAD1 T726 at unattached kinetochores (Figure 3D,E), which is critical for MCC (mitotic checkpoint complex) assembly and consequently for a functional checkpoint. Therefore, we conclude that the phosphorylation of PP1α87B at T286 by POLO is required for efficient MPS1 activity at unattached kinetochores and robust SAC signaling.

**Figure 3.**
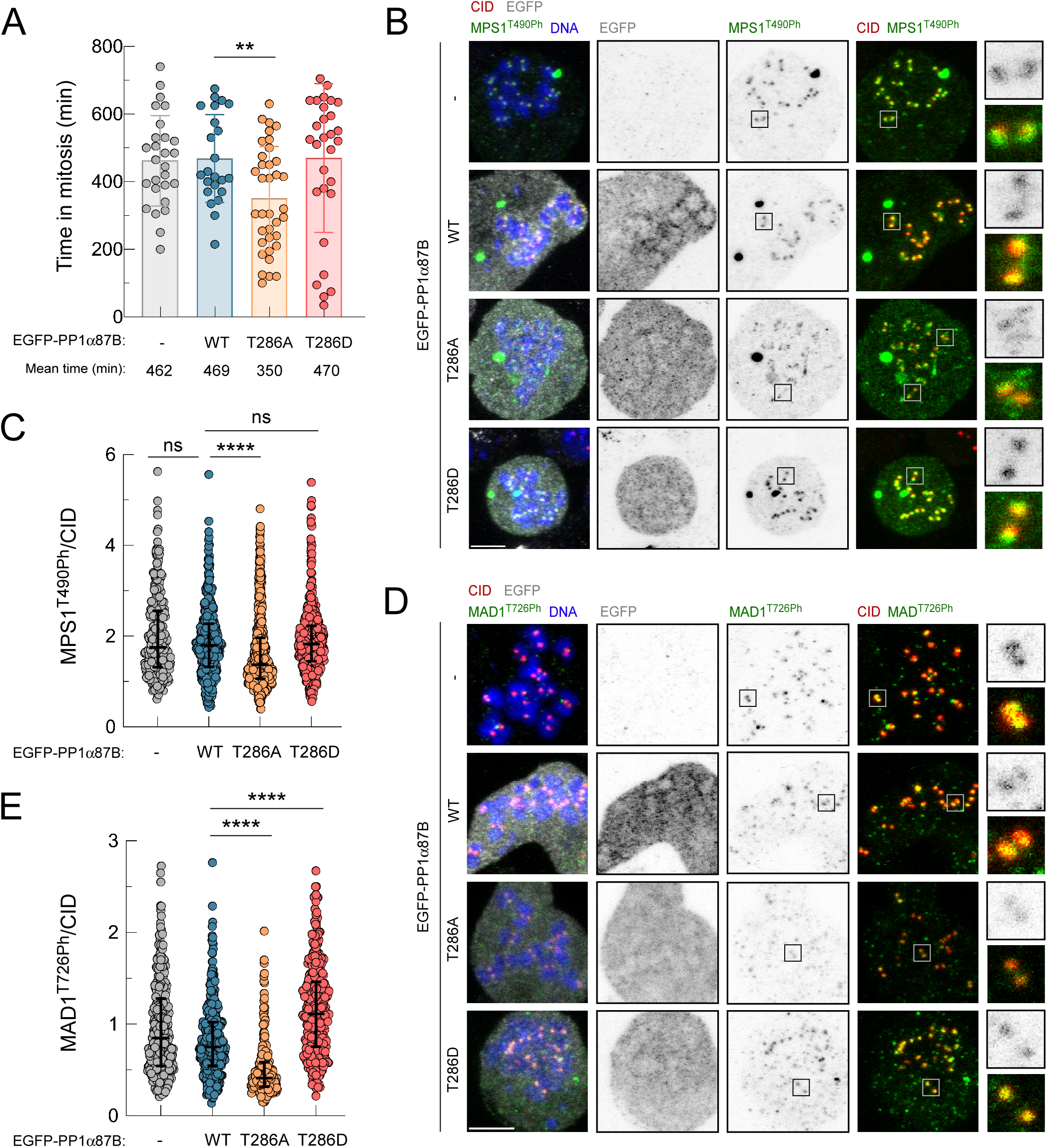
Phosphorylation of PP1α87B at T286 is required for MPS1 activation at unattached kinetochores and robust SAC signaling. (A) Mitotic duration of *Drosophila* S2 cells expressing the indicated EGFP-PP1α87B transgenes in the presence of colchicine. The time in mitosis was monitored by time-lapse microscopy and corresponds to the length of time between the nuclear envelope breakdown and mitotic exit, identified by chromatin decondensation (n ≥ 23 cells for each condition). (B,C) Representative immunofluorescence images (B) and corresponding quantifications (C) of MPS1 T490 phosphorylation (MPS1^T490Ph^) levels at kinetochores of S2 cells expressing the indicated transgenes in the presence of colchicine for 6 hours. MPS1^T490h^ fluorescence intensity was determined relative to CID signal (n ≥ 284 kinetochores from at least 20 cells for each condition). (D,E) Representative immunofluorescence images (D) and corresponding quantifications (E) of MAD1 T726 phosphorylation (MAD1^T726Ph^) levels at kinetochores of S2 cells expressing the indicated transgenes in the presence of colchicine for 6 hours. MAD1^T726Ph^ fluorescence intensity was determined relative to CID signal (n ≥ 242 kinetochores from at least 15 cells for each condition). Data information: in A, data are presented as mean with standard deviation; in C and E, data are presented as median with interquartile range; asterisks indicate that differences between mean ranks are statistically significant, **p<0.01; ****p<0.0001; ns, non-significant (Kruskal-Wallis, Dunn’s multiple comparison test in C, E and Mann-Whitney test in A). Scale bars: 5μm.

### Mimicking constitutive phosphorylation of PP1α87B at T286 prevents timely SAC silencing

The above results imply that phosphorylation of PP1α87B at T286 attenuates its ability to dephosphorylate the activating T-loop of MPS1 recruited to unattached kinetochores and thereby strengthens the SAC. Hence, we next assessed the impact of mimicking permanent phosphorylation of T286 (T286D) throughout mitosis. We first evaluated the activation of MPS1 in prometaphase cells. As expected, immunofluorescence analysis demonstrated enhanced phosphorylation of the MPS1 T-loop at prometaphase kinetochores in PP1α87B depleted cells (Figure S3A-C). This effect was rescued by the expression of EGFP-PP1α87B^WT^ but less efficiently by EGFP-PP1α87B^T286D^ (Figure S3A-C). Importantly, kinetochore levels of MPS1^T490Ph^ were significantly reduced in EGFP-PP1α87B^T286A^-expressing cells as compared to control parental cells or cells expressing the wild type transgene (Figure S3A-C). These results confirm that preventing the phosphorylation of PP1α87B T286 renders MPS1 less active at prometaphase kinetochores, as observed for fully unattached kinetochores in colchicine-treated cells.

We subsequently assessed the activation status of MPS1 at metaphase kinetochores. Immunofluorescence analysis revealed that the expression of EGFP-PP1α87B^WT^ or EGFP-PP1α87B^T286A^ rescued the increase of active MPS1 (MPS1^T490Ph^) caused by the depletion of endogenous PP1α87B (Figure 4A-C). In contrast, cells expressing the phosphomimetic version EGFP-PP1α87B^T286D^ failed to efficiently dephosphorylate MPS1 T-loop (Figure 4A-C), even though inter-kinetochore distances were similar to those measured in bioriented chromosomes of control parental or EGFP-PP1α87B^WT^ cells and compatible with stable kinetochore-microtubule attachments (Figure 4D,E). In line with the inability to inactivate MPS1, EGFP-PP1α87B^T286D^-expressing cells were delayed in the transition to anaphase when compared to cells expressing EGFP-PP1α87B^WT^ or EGFP-PP1α87B^T286A^, which fully rescued the prolonged metaphase duration caused by the depletion of endogenous PP1α87B (Figure 4F). Based on these results, we conclude that mimicking PP1α87B constitutively phosphorylated at T286 impairs timely inactivation of MPS1 and thereby delays SAC silencing and anaphase onset, despite chromosome biorientation and normal inter-kinetochore tension.

**Figure 4.**
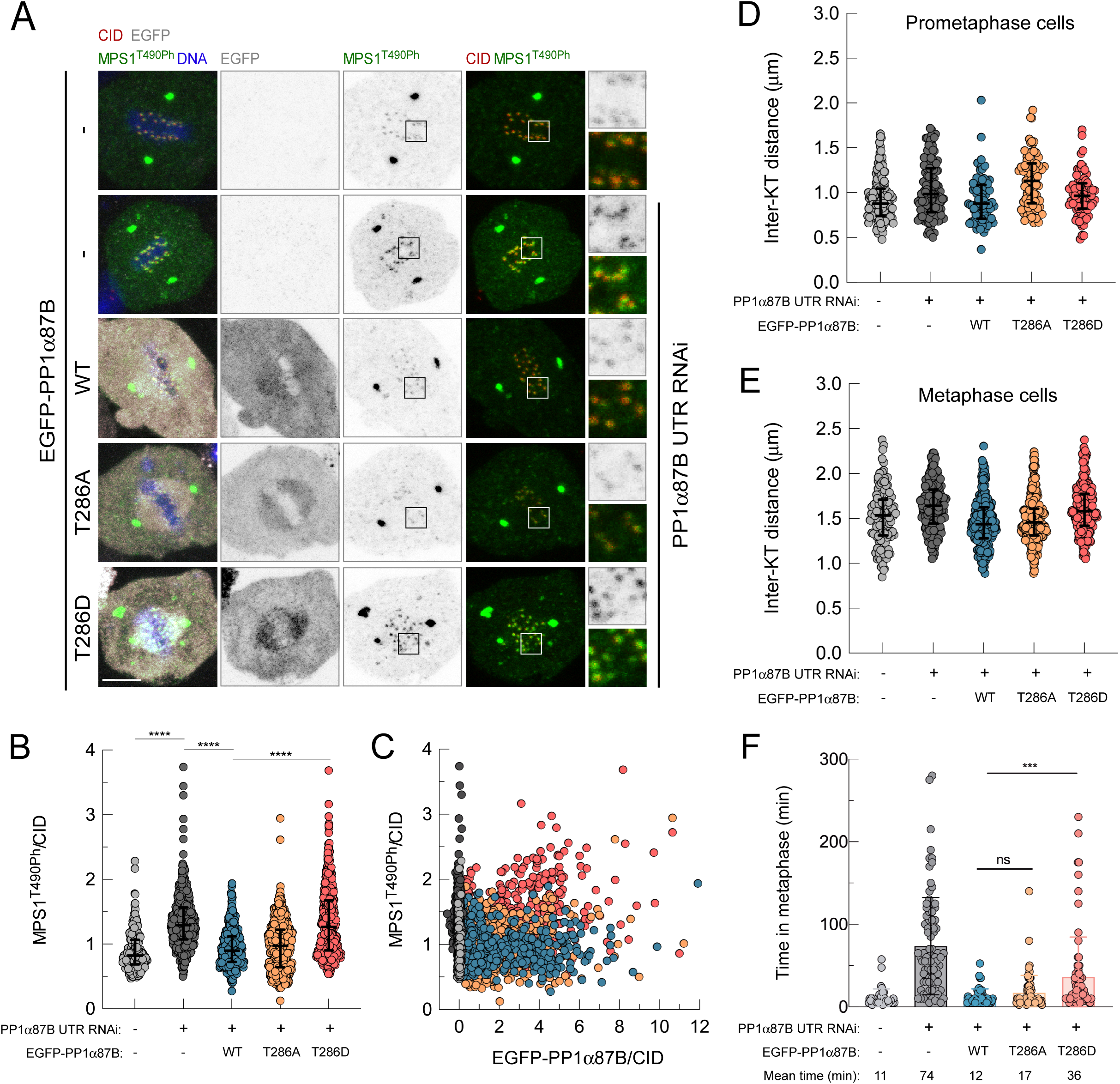
Mimicking constitutive phosphorylation of PP1α87B T286 prevents timely MPS1 inactivation and SAC silencing in metaphase. (A-C) Representative immunofluorescence images (A) and corresponding quantifications (B) of MPS1 T490 phosphorylation (MPS1^T490Ph^) levels at kinetochores of control and PP1α87B-depleted S2 cells in metaphase expressing the indicated EGFP-PP1α87B transgenes. (C) MPS1^T490Ph^ levels in B plotted over EGFP-PP1α87B levels at kinetochores. MPS1^T490h^ and EGFP fluorescence intensities were determined relative to CID signal (n ≥ 229 kinetochores for each condition). (D,E) Quantifications of inter-kinetochore distances of control and PP1α87B-depleted S2 cells in prometaphase (D) and in metaphase (E) expressing the indicated EGFP-PP1α87B transgenes. Inter-kinetochore distance was measured as the distance between centroids of CID pairs (n ≥ 77 sister-kinetochore pairs for each condition). (F) Metaphase duration of control and PP1α87B-depleted S2 cells expressing the indicated EGFP-PP1α87B transgenes. The time in metaphase was monitored by time-lapse microscopy and corresponds to the length of time measured between the frame when all chromosomes align at the metaphase plate and the frame when anaphase onset occurs (n ≥ 40 cells for each condition). Data information: data in B, D and E are presented as median with interquartile range and data in F are presented as mean with standard deviation; asterisks indicate that differences between mean ranks are statistically significant, ***p<0.001; ****p<0.0001; ns, non-significant (Kruskal-Wallis, Dunn’s multiple comparison test). Scale bars: 5μm. See also Figure S3.

### Kinetochore-tethering does not rescue the inhibition exerted by T286 phosphorylation on the SAC-antagonizing activity of PP1α87B

The recruitment of PP1α87B to kinetochores is required for SAC satisfaction in *Drosophila* S2 cells (Audett et al., 2022). However, forcing kinetochore accumulation of a wild type version of PP1α87B by fusing it to MIS12 did not cause premature inactivation of the SAC (Figure 1A). This suggests that the weakened SAC observed in EGFP-PP1α87B^T286A^ mutants and the delayed SAC silencing in EGFP-PP1α87B^T286D^ cells are unlikely due to excessive or deficient kinetochore recruitment of PP1α87B, respectively. To directly delineate the effect of T286 phosphorylation on the activity of PP1α87B, we bypassed potential influences from kinetochore recruitment or dynamics by tethering the T286 phosphomutants to MIS12 in cells depleted of endogenous PP1α87B. This ensures consistent phosphatase abundance and behavior at the outer kinetochore for direct comparison. Unlike MIS12-EGFP-PP1α87B^WT^ and MIS12-EGFP-PP1α87B^T286A^, cells expressing MIS12-EGFP-PP1α87B^T286D^ failed to efficiently dephosphorylate the T-loop of MPS1 and retained elevated levels of MAD1^T726Ph^ at metaphase kinetochores (Figure S4A-C). Accordingly, MIS12-EGFP-PP1α87B^T286D^ cells exhibited a prolonged metaphase delay, whereas expression of MIS12-EGFP-PP1α87B^WT^ or MIS12-EGFP-PP1α87B^T286A^ restored timely anaphase onset in a PP1α87B RNAi background (Figure S4D). The metaphase delay observed in MIS12-EGFP-PP1α87B^T286D^ was SAC-dependent, as it was averted by MPS1 depletion (Figure S4D). Collectively, these results with the kinetochore-tethered PP1α87B phosphomutants strongly suggest that phosphorylation of T286 inhibits PP1α87B-mediated SAC silencing through mechanisms other than a possible impact on PP1α87B kinetochore recruitment or dynamics.

### Phosphorylation of PP1α87B at T286 cooperates with limited kinetochore recruitment of PP1α87B to ensure robust SAC signaling at unattached kinetochores

Next, we evaluated the impact of the kinetochore-tethered PP1α87B phosphomutants on SAC strength by monitoring their ability to delay mitotic exit in the presence of colchicine (Figure 5A). Similarly to the untethered counterparts, expression of MIS12-EGFP-PP1α87B^WT^ and MIS12-EGFP-PP1α87B^T286D^ enabled cells to arrest in mitosis. This indicates that SAC proficiency was retained despite forced phosphatase accumulation at unattached kinetochores. On the other hand, preventing T286 phosphorylation of the kinetochore-tethered phosphatase (MIS12-EGFP-PP1α87B^T286A^) significantly accelerated mitotic exit, which is symptomatic of a compromised SAC. Notably, the SAC was substantially weaker than in cells expressing untethered EGFP-PP1α87B^T286A^ (∼165 min vs ∼350 min of mitotic arrest duration, respectively). This indicates that increasing the kinetochore localization of PP1α87B can drive SAC inactivation at unattached kinetochores, as long as T286 is not phosphorylated. Accordingly, a stronger reduction in the levels of MPS1^T490Ph^ at unattached kinetochores was observed in colchicine-treated cells expressing MIS12-EGFP-PP1α87B^T286A^ (50% reduction relative to parental cells, Figure 5B,C) than in those expressing EGFP-PP1α87B^T286A^ (25% reduction relative to parental cells, Figure 3B,C). These results suggest that phosphorylation of PP1α87B at T286 by POLO cooperates with AURORA B-mediated control of PP1α87B recruitment to the kinetochore to efficiently limit phosphatase activity at unattached kinetochores and ensure SAC proficiency.

**Figure 5.**
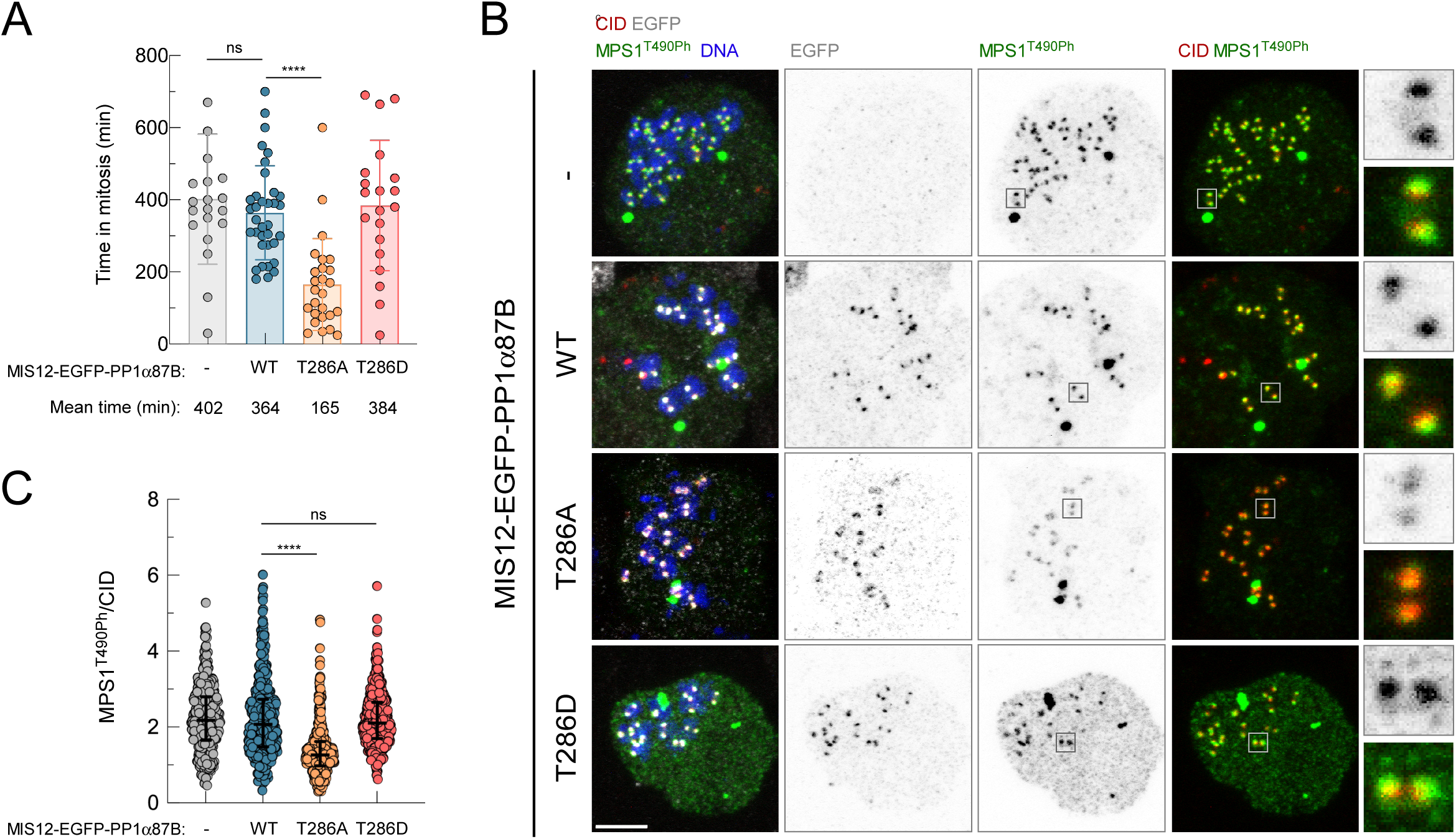
Rendering kinetochore-tethered PP1α87B unphosphorylatable at T286 compromises MPS1 activation and SAC signaling in colchicine-incubated S2 cells. (A) Mitotic duration of *Drosophila* S2 cells expressing the indicated MIS12-EGFP-PP1α87B transgenes in the presence of colchicine. The time in mitosis was monitored by time-lapse microscopy and corresponds to the length of time between the nuclear envelope breakdown and mitotic exit, identified by chromatin decondensation (n ≥ 20 cells for each condition). (B,C) Representative immunofluorescence images (B) and corresponding quantifications (C) of MPS1 T490 phosphorylation (MPS1^T490Ph^) levels at kinetochores of S2 cells expressing the indicated MIS12-EGFP-PP1α87B transgenes in the presence of colchicine for 6 hours. MPS1^T490h^ fluorescence intensity was determined relative to CID signal (n ≥ 392 kinetochores for each condition). Data information: data in A are presented as mean with standard deviation and data in C are presented as median with interquartile range; asterisks indicate that differences between mean ranks are statistically significant, ****p<0.0001; ns, non-significant (Kruskal-Wallis, Dunn’s multiple comparison test). Scale bars: 5μm. See also Figure S4.

### Phosphorylation of PP1α87B/PP1 at T286/T288 does not directly affect its catalytic activity in vitro

Next, we set out to dissect how T286 phosphorylation by POLO opposes the ability of PP1α87B to dephosphorylate and inactivate kinetochore-localized MPS1. To test if phosphorylation of T286 in *Drosophila* PP1α87B or T288 in human PP1α/γ directly inhibits their catalytic activity, we purified bacterially-produced wild type and phosphomutant versions of these phosphatases and assessed their capacity to dephosphorylate 8-difluoro-4-methylumbelliferone phosphate (DiFMUP), an artificial fluorogenic substrate lacking PP1-interacting SLiMs. We did not observe significant changes in DiFMUP dephosphorylation rate between the wild type and the phosphomimetic or phosphodefective versions of the phosphatases (Figure S5A-C). Neither did we detect an impact on the activity of wild type PP1γ by PLK1-mediated phosphorylation at T288 (Figure S5C). Conversely, all phosphatases were consistently inhibited by okadaic acid, a well-established inhibitor of PP1 and PP2A phosphatases (Garcia et al., 2002), thereby confirming that the purified recombinant proteins used in the assay were active (Figure S5A-C). These results show that the *in vitro* enzymatic activity of PP1 is not affected by its phosphorylation at T286/T288. This is consistent with the lack of noticeable structural changes or catalytic impairments in PP1α87B after conversion of T286 to aspartate (T286D) or alanine (T286A) as predicted by the Colabfold variant of AlphaFold 2 (Jumper et al., 2021; Mirdita et al., 2022) (Figure S5D,E).

### Phosphorylation of PP1α87B at T286 decreases its interaction with MPS1

Interestingly, T286/T288 is located on a small surface loop distant from the active center (Figure 2B, Figure S5D,E) but positioned at the entrance of the PP1α87B/PP1γ hydrophobic pocket that binds to RVxF motifs, directly contributing to the interaction in some cases (Ragusa et al., 2010; Fedoryshchak et al., 2020). This prompted us to investigate whether phosphorylation of PP1α87B/PP1γ at T286/T288 reduces its binding to *Drosophila* MPS1, which has an RVxF-type PP1-binding SLiM (^231^KVLF^234^) near the N-terminus. (Moura et al., 2017). Pull-down experiments with purified *Drosophila* MBP-MPS1^104-330^ and 6xHis-PP1γ^WT^ confirmed their interaction (Figure 6A), which was, however, significantly compromised after alanine mutation of the RVxF motif (^231^AAAA^234^) of MPS1 or by converting the T288 of PP1γ into aspartate (T288D) (Figure 6A). This is consistent with AlphaFold 3 modeling of PP1α87B binding to the MPS1 ^231^KVLF^234^, with the latter being dislodged from the hydrophobic groove of PP1α87B when T286 is phosphorylated (Figure 6B, Figure S5F). These results suggest that T286 phosphorylation hinders PP1α87B-mediated inactivation of MPS1 due to defective binding to the kinase N-terminus. To validate this hypothesis in a cellular context, we artificially fused PP1α87B^T286D^ to MPS1 and assessed its capacity to antagonize MPS1 T-loop phosphorylation at metaphase kinetochores. Through time-lapse imaging we first confirmed that inducing EGFP-MPS1 overexpression in S2 cells caused a prolonged mitotic arrest (Figure 6C), which correlates with elevated levels of active MPS1 on metaphase kinetochores (Figure 6D-F), as previously reported (Conde et al., 2013). Tethering PP1α87B^WT^ to EGFP-MPS1 efficiently attenuated the increase in MPS1 T-loop phosphorylation (Figure 6D-F) and significantly reduced the delay in mitosis (Figure 6C). Importantly, PP1α87B^T286D^ fused to EGFP-MPS1 was equally effective in counteracting MPS1 activation at kinetochores and accordingly decreased the mitotic duration to the same extent as observed in cells overexpressing PP1α87B^WT^-EGFP-MPS1 (Figure 6C-F). Conversely, fusing EGFP-MPS1 with a catalytic inactive version of the phosphatase (PP1α87B^D62N^-EGFP-MPS1) failed to ameliorate the mitotic arrest driven by excessive MPS1 activity (Figure 6C-F). These results show that artificially forcing PP1α87B binding to MPS1 in S2 cells bypasses the inhibition that T286 phosphorylation exerts on PP1α87B-mediated inactivation of MPS1. In conclusion, our *in silico*, *in vitro* and *in cellulo* approaches support a molecular mechanism whereby phosphorylation of T286 hinders the ability of PP1α87B hydrophobic pocket to engage with the RVxF motif of MPS1, thereby precluding the binding of PP1α87B to MPS1 and ultimately impairing dephosphorylation of the MPS1 T-loop and SAC silencing.

**Figure 6.**
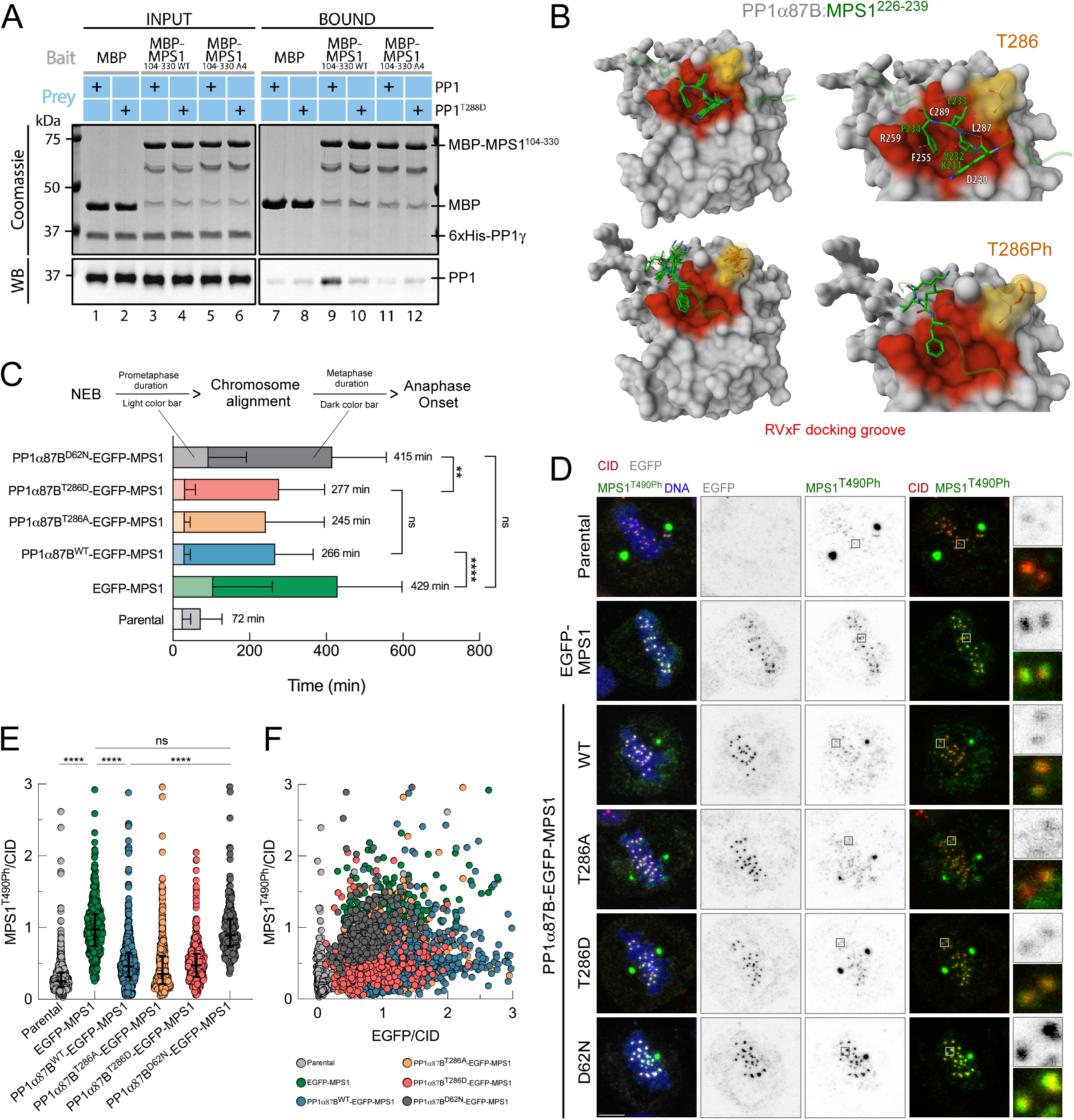
Phosphorylation of PP1α87B/PP1γ T286/T288 opposes direct binding of PP1α87B/PP1γ to MPS1. (A) Pull-downs of human 6xHis-PP1γ^WT^ and 6xHis-PP1γ^T288D^ by bead-immobilized *Drosophila* MBP-MPS1^104-330^ ^WT^ (^231^KVLF^234^) and MBP-MPS1^104-330^ ^A4^ (^231^AAAA^234^). All indicated recombinant proteins were produced in and purified from bacteria. Input and beads (bound) were resolved by SDS-PAGE, stained with coomassie and blotted for 6xHis-PP1γ. (B) AlphaFold 3 modelling of the interaction between the RVxF motif (^231^KVLF^234^) of *Drosophila* MPS1 (Q9VEH1) and PP1α87B (P12982). The structural representation on the upper left shows the superimposition of 5 models with ^231^KVLF^234^ residues (labelled in green) of the MPS1^226-239^ peptide perfectly docked into the RVxF-binding pocket (highlighted in red) of PP1α87B. A magnification of the interaction is shown on the upper right representation with the main interacting residues from the RVxF-binding pocket labelled in white and unphosphorylated T286 (highlighted in yellow) indicated. Predicted hydrogen bonds are represented as a dashed blue cylinder and π-π bonds are represented as a dashed lime cylinder. The lower left structural representation shows the superimposition of 5 models predicting that T286 phosphorylation (T286Ph) dislodges the KVLF of MPS1 from the RVxF-binding pocket. The corresponding magnification of the RVxF-binding groove is presented on the lower right. Magnifications display rank 1 of 5 modeling predictions for each complex. (C) Mitotic timing of S2 cells overexpressing the indicated transgenes. Light-colored bars represent the prometaphase duration (length of time measured between nuclear envelope breakdown, NEB, and the first frame with all chromosomes aligned at the metaphase plate; n ≥ 20 cells for each condition) and dark-colored bars represent the metaphase duration (length of time measured between the first frame after chromosome alignment and anaphase onset, n ≥ 23 cells for each condition). (D-F) Representative immunofluorescence images (D) and corresponding quantifications (E,F) of MPS1 T490 phosphorylation (MPS1^T490Ph^) levels at kinetochores of metaphase S2 cells overexpressing the indicated transgenes. MPS1^T490h^ and EGFP fluorescence intensities were determined relative to the CID signal (n ≥ 259 kinetochores for each condition). Data information: data in C are presented as mean with standard deviation; asterisks indicate that differences between means are statistically significant, ****p<0.0001; ns, non-significant (Two-Way ANOVA, Tukey’s multiple comparison test). Data in E are presented as median with interquartile range; asterisks indicate that differences between mean ranks are statistically significant, ****p<0.0001; ns, non-significant (Kruskal-Wallis, Dunn’s multiple comparison test). Scale bars: 5μm. See also Figure S5.

### Phospho-regulation of PP1α87B by POLO is required for genome stability in vivo

The results obtained with cultured S2 cells demonstrate that POLO phosphorylates PP1α87B at T286 during early mitosis to limit the dephosphorylation of kinetochore-associated MPS1 and ensure robust SAC signaling. Conversely, the phosphorylation of PP1α87B^T286^ at kinetochores concomitantly decreases with chromosome biorientation to enable timely SAC silencing and anaphase onset. To investigate whether this phospho-regulation occurs *in vivo* and its relevance for genome stability, we analyzed the impact of PP1α87B^T286^ phosphomutants on dividing neuroblasts of 3rd instar larvae. We used inscuteable-GAL4/UAS to drive the expression of PP1α87B-HA transgenes in a *pp1α87B^1^/pp1α87B^87Bg-3^* trans-heterozygous genetic background (hypomorphic/amorphic heteroallelic combination), estimated to have residual endogenous activity of total PP1 (∼20%) in 3rd instar larvae (Kirchner et al., 2007). MPS1 activation was assessed upon colchicine treatment and, as in cultured S2 cells, reducing endogenous PP1α87B activity led to an increase of MPS1 T-loop phosphorylation at unattached kinetochores, which was efficiently prevented by the expression of PP1α87B^WT^-HA (Figure 7A,B). Unphosphorylatable PP1α87B^T286A^-HA reduced the kinetochore levels of MPS1^T490Ph^ even further, whereas phosphomimetic PP1α87B^T286D^-HA failed to rescue MPS1 hyper-activation (Figure 7A,B). We then monitored by live imaging neuroblasts undergoing anaphase to evaluate the accuracy of genome partitioning (Videos 3-7). Approximately 55% of *pp1α87B^1^/pp1α87B^87Bg-3^* trans-heterozygous mutant neuroblasts displayed chromosome segregation defects, which were largely averted by expression of PP1α87B^WT^-HA (Figure 7C,D). However, both phosphomutants failed to restore the fidelity of chromosome segregation as efficiently as the wild type transgene, with PP1α87B^T286A^-HA neuroblasts exhibiting lagging chromosomes in approximately 27% of anaphases and asynchronous segregation of chromosomes being scored in 25% of dividing PP1α87B^T286D^-HA neuroblasts (Figure 7C,D). In line with the observed segregation defects, the percentage of aneuploid neuroblasts in the larval brains increased significantly when the phospho-regulation of PP1α87B^T286^ was compromised (Figure 7E,F). Collectively, these results strongly support the notion that timely (de)phosphorylation of PP1α87B at T286 is critical for faithful chromosome segregation, and consequently, for genome stability *in vivo*. In line with the physiological importance of tightly regulating PP1α87B^T286^ phosphorylation, ubiquitous expression of either UAS-PP1α87B-HA phosphomutant was less efficient in rescuing the lethality of *pp1α87B^1^/pp1α87B^87Bg-3^* trans-heterozygotes than the expression of UAS-PP1α87B^WT^-HA (Figure 7G,H). It is important to note that the ubiquitous expression of PP1α87B^WT^-HA could not fully complement the *pp1α87B^1^/pp1α87B^87Bg-3^* mutant background (Figure 7G,H). Hence, we cannot exclude that arm-GAL4-dependent expression of UAS-PP1α87B results in protein levels that are insufficient to sustain a complete rescue in viability of *pp1α87B^1^/pp1α87B^87Bg-3^* mutants. Nevertheless, the results show a consistent inability of both PP1α87B phosphomutants to restore viability to the same extent as the wild type PP1α87B, underscoring the physiological importance of its regulation by T286 phosphorylation (Figure 7G,H).

**Figure 7.**
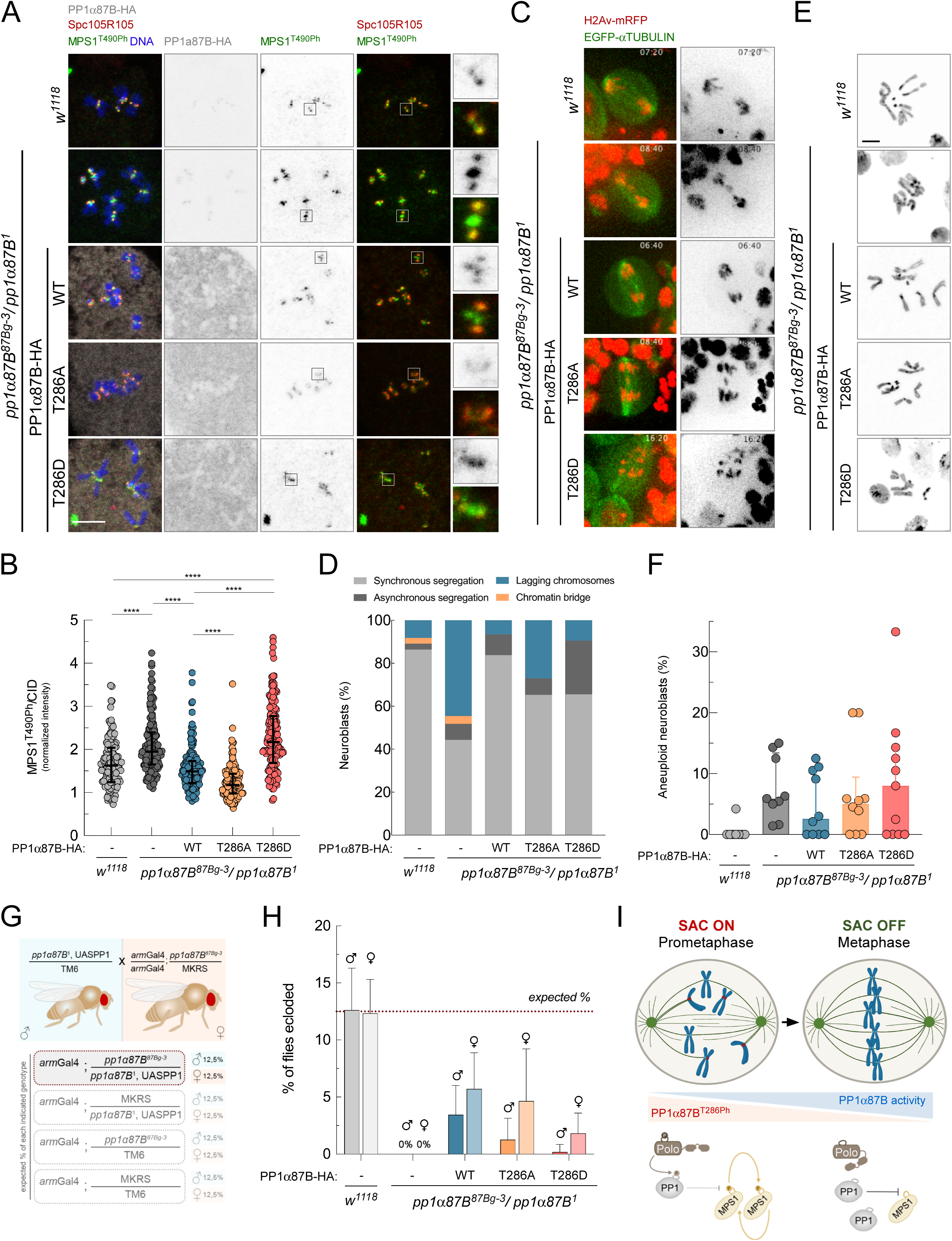
Reversible phosphorylation of PP1α87B T286 is required for mitotic fidelity and genome stability of *Drosophila* neuroblasts. (A,B) Representative immunofluorescence images (A) and corresponding quantifications (B) of MPS1 T490 phosphorylation (MPS1^T490h^) levels at kinetochores of colchicine-incubated neuroblasts from 3rd instar larvae with the indicated genotypes. MPS1^T490Ph^ fluorescence intensities were determined relative to SPC105R signal (n ≥ 121 kinetochores for each condition). (C,D) Representative video stills (C) of neuroblasts from 3rd instar larvae with the indicated genotypes undergoing anaphase and corresponding quantifications (D) of the frequency of neuroblasts with the indicated chromosome segregation anomalies. The mitotic progression of neuroblasts was monitored by time-lapse microscopy (n ≥ 26 neuroblasts from each genotype). (E, F) Representative immunofluorescence images (E) of mitotic neuroblasts from squashed brains showing distinct chromosome content and corresponding quantification (F) of the percentage of aneuploid neuroblasts in 3rd instar larvae with the indicated genotypes. For the quantification of aneuploidy, the whole brain was analysed and prometaphase and metaphase neuroblasts were scored for chromosome number. An aneuploid cell was considered when more or less than 8 chromosomes were visualized. Each dot represents an independent brain (n ≥ 184 neuroblasts from at least 8 larvae brains of each genotype). (G) Schematic representation of the genetic crosses performed for the rescue experiment of *pp1α87B^B1^/pp1α87B^87Bg-3^*trans-heterozygous lethal phenotype. The expected percentage of each possible genotype in F1 is indicated for both males and females (M&M for details). (H) Quantification of the percentage of F1 males and females with the genotype highlighted in G. Control crosses were performed using males carrying UASLacZ and females carrying armGal4 (n ≥ 6 independent crosses for each condition). (I) Proposed model for the control of SAC signaling by POLO-mediated phosphorylation of PP1α87B. Phosphorylation of T286 at prometaphase kinetochores limits PP1α87B interaction with MPS1 thereby leaving the activating T-loop autophosphorylation of MPS1 unopposed to ensure a robust SAC. POLO activity decreases at microtubule-attached kinetochores in metaphase, which together with dephosphorylation of T286, license PP1α87B for MPS1 binding and T-loop dephosphorylation. This inactivates MPS1 and consequently silences the checkpoint signaling pathway to allow timely anaphase onset. Data information: data in B are presented as median with interquartile range, data in D are plotted as fractions of the total, data in F are presented as median with interquartile range and data in H are presented as mean with standard deviation; asterisks indicate that differences between mean ranks are statistically significant, ****p<0.0001; ns, non-significant (Kruskal-Wallis, Dunn’s multiple comparison test). Scale bars: 5μm.

## DISCUSSION

PP1 catalyzes a major fraction of all dephosphorylation events in eukaryotic cells and is crucial for cell cycle progression (Moorhead et al., 2007). By switching-off the upstream SAC orchestrator (MPS1) and by antagonizing its downstream phosphorylations, PP1 ensures prompt SAC silencing at kinetochores following chromosome biorientation and thereby avoids unnecessary and potentially genome-menacing delays in mitosis (Orth et al., 2012; Shindo et al., 2021; Jain and Tran, 2023). Despite this well-established requirement of PP1 for timely anaphase onset and mitotic exit, a coherent model of how PP1 is regulated during mitosis was still missing. It had remained unclear how PP1 activity is repressed during prometaphase to ensure robust SAC function but becomes increasingly active in metaphase to drive SAC silencing. Here, we provide novel insights into the mitotic regulation of PP1. We identify a previously uncharacterized mechanism by which POLO modulates PP1α87B function (Figure 7I). Specifically, we demonstrate that POLO phosphorylates PP1α87B at a conserved residue (T286) within its RVxF-docking groove, hindering its binding to the RVxF motif of MPS1 and consequently preventing the dephosphorylation of the MPS1 T-loop. Phosphorylation of PP1α87B on T286 is most pronounced at unattached or tensionless kinetochores, which is in line with the elevated kinetochore levels of active POLO and the need for robust SAC signaling. In this context, AURORA B-mediated phosphorylation of SILK and RVxF motifs on SPC105R (KNL1 ortholog) limits the recruitment of PP1α87B to the outer kinetochore. However, forcing outer kinetochore accumulation of PP1α87B at unattached kinetochores only marginally contributed to inactivating MPS1 and antagonizing the SAC. In contrast, rendering PP1α87B^T286^ unphosphorylatable (T286A) reduced MPS1 activation at unattached kinetochores (decreased MPS1 T-loop phosphorylation), which, accordingly, deterred MPS1 activity towards SAC substrates such as MAD1^T726^ and precluded robust SAC signaling (decreased mitotic arrest in the presence of colchicine). MPS1 inactivation and weakening of the SAC were even more pronounced when the PP1α87B^T286A^ variant was tethered to unattached kinetochores, indicating that AURORA B-dependent reduction in the recruitment of PP1α87B to kinetochores synergizes with T286 phosphorylation to efficiently dampen the activity of PP1α87B at unattached/prometaphase kinetochores and ensure a robust SAC. Accordingly, mimicking constitutive phosphorylation of T286 (T286D) precludes efficient MPS1 inactivation at metaphase kinetochores and concomitantly delays SAC silencing and anaphase onset. Hence, dephosphorylation of T286 is required for PP1α87B to become sufficiently active in metaphase, rendering the SAC responsive to end-on microtubule attachments and ensuring a timely cell cycle progression. The molecular underpinnings of T286 dephosphorylation are presently elusive but merit further investigations. Even minor delays in metaphase can lead to precocious and insufficient activation of SEPARASE, which results in chromosome nondisjunction in anaphase due to incomplete COHESIN removal (Shindo et al., 2021). Accordingly, we found that fine-tuning PP1α87B activity by POLO is required for mitotic fidelity and genome stability *in vivo*. Specifically interfering with T286 phosphorylation in dividing *Drosophila* neuroblasts led to chromosome segregation errors and to a concurrent accumulation of aneuploid cells in larval brains. Neuroblasts expressing unphosphorylatable PP1α87B^T286A^ often exhibited anaphases with lagging chromosomes, whereas expression of PP1α87B^T286D^ caused asynchronous migration of segregating chromosomes towards the spindle poles. These phenotypes are consistent with an inhibitory role of T286 phosphorylation on PP1α87B activity. Inefficient repression of PP1α87B activity in the absence of T286 phosphorylation (T286A) precludes robust MPS1 activity. This not only weakens the SAC but also hampers pole-based correction of erroneous kinetochore-microtubule attachments in flies (Leça et al., 2024 *Preprint*). As a result, PP1α87B^T286A^ neuroblasts undergo anaphase before chromosome biorientation, leading to chromosome mis-segregation and aneuploidy. Conversely, constitutively inhibited PP1α87B in PP1α87B^T286D^-expressing neuroblasts prevents timely MPS1 inactivation and SAC silencing. This prolongs the metaphase duration, leading to precocious SEPARASE activity and asynchronous chromosome segregation in the ensuing anaphase (Shindo et al., 2021). These data demonstrate that fine-tuning PP1α87B activity through timely (de)phosphorylation of T286 is critical for SAC robustness and responsiveness *in vivo* and underscores the requirement of these checkpoint properties for genome stability in a proliferative tissue.

Whether phosphorylation of PP1γ at T288 by PLK1 also impairs SAC silencing in human cells is presently unclear, but would be consistent with the observation that PLK1 removal from kinetochores is a crucial event in SAC silencing (Cordeiro et al., 2020). However, human MPS1 lacks a canonical RVxF motif, and, accordingly, PP1 seems to be dispensable for MPS1 inactivation in HeLa cells (Hayward et al., 2019). Instead, end-on microtubule attachments displace a significant fraction of MPS1 from kinetochores, thereby limiting its activity at kinetochores and consequently SAC signaling. Hence, in mammalian cells, PLK1-mediated regulation of PP1γ activity is expected to modulate the SAC signaling pathway downstream of MPS1. Future efforts to elucidate if inhibition of PP1 by PLK1 is a conserved feature of SAC signaling and to identify which substrates fail to bind to PP1γ when T288 is phosphorylated are warranted.

The recruitment of POLO-kinase family members to different locations and substrates is typically mediated by a conserved POLO-Box domain (PDB) that binds to phosphorylated S-pS/pT-X motifs (Elia et al., 2003). In many cases, X is a proline, and this S-pS/pT-P motif constitutes a CDK-primed POLO binding motif. However, PP1α87B lacks an obvious POLO binding motif. This suggests that in a cellular context, POLO either directly binds to PP1α87B through a non-canonical mechanism or relies on an intermediary protein to bridge the interaction. Examples of non-canonical POLO/PLK1 interactions occur with BORA (Seki et al., 2008), MKLP2 (Neef et al., 2003), MTRM (Bonner et al., 2013), and MAP205 (Archambault et al., 2008). It is also possible that POLO targets PP1α87B when docked onto other proteins, as in the case of PLK1 bound to the BUB complex, which directly phosphorylates KNL1 MELT motifs in human cells and *C. elegans* (Espeut et al., 2015; von Schubert et al., 2015; Ikeda and Tanaka, 2017; Cordeiro et al., 2020). In doing so, BUB1-bound PLK1 primes its own recruitment and locally amplifies SAC signaling by boosting KNL1 phosphorylation independently of MPS1 (Cordeiro et al., 2020). It will be important to test if the same pool of PLK1 also phosphorylates KNL1-bound PP1, and if so, whether it contributes to further reinforce the KNL1-BUB-PLK1 positive feedback loop at unattached kinetochores. Additional proteins bridging a POLO/PLK1-PP1α87B/PP1 interaction are MYPT1 and APOLO1, which have been characterized as RIPPOs that fine-tune PLK1 activity at kinetochores (Yamashiro et al., 2008; Dumitru et al., 2017; Xu et al., 2021). Whether PLK1 can also directly modulate the phosphatase activity of PP1 in these complexes is not known. Identifying bridging proteins, the pools of PLK1 involved, and the subset of PP1 holoenzymes that are targeted are key steps in further elucidating the molecular mechanisms underlying PP1α87B/PP1 regulation by POLO/PLK1.

Previous studies have primarily focused on the phosphorylation of RVxF motifs themselves as a means of regulating PP1 binding and activity towards substrates. In the RVxF motif, x is any residue and V/F constitute the principal PP1-binding residues. Position x is a S or T residue in 21% and 18% of the variants, respectively, and their phosphorylation, mainly catalyzed by AURORA B in mitosis, reduces the affinity for PP1 (Dent et al., 1990; Liu et al., 2010; Nasa et al., 2018). We now demonstrate that the RVxF-binding groove on the PP1 catalytic subunit is itself phosphorylated to achieve a similar purpose. We speculate that this regulatory mechanism governed by POLO/PLK1 evolved to control PP1α87B/PP1 binding to RVxF motifs that lack a phosphorylatable S/T at the “x” position, as showcased by the KVLF of MPS1. This ensures that PP1α87B/PP1 recruitment to AURORA B-impervious RIPPOs can still be regulated in a phosphorylation-dependent manner in response to specific cellular conditions, such as kinetochore-microtubule attachment status. By employing two independent, but not necessarily exclusive, regulatory mechanisms orchestrated by distinct kinases (POLO/PLK1 and AURORA B), cells can achieve highly dynamic, versatile, graded and responsive thresholds of PP1α87B/PP1 activity across mitotic structures. This nuanced control of PP1α87B/PP1 enables the dephosphorylation of different substrates in specific subcellular compartments and at various stages of mitotic progression, thereby ensuring precise control of mitotic events so that these occur in the correct order and at the appropriate time.

Our data argue against the notion that CDK1-mediated phosphorylation of the PP1 C-terminal tail plays a significant role in regulating SAC-antagonizing PP1 activity, at least in *Drosophila* cells. This is consistent with the observation that efficient APC/C-triggered proteolysis of CYCLIN B is contingent upon SAC inactivation, a process that itself requires PP1 activity. However, our study does not exclude that the phosphorylation of the C-terminal tail of PP1 serves to inhibit a subset of PP1 holoenzymes involved in other mitotic and cell cycle events.

In summary, our findings expand the repertoire of regulatory strategies that cells employ to control PP1α87B/PP1 activity in mitosis. By phosphorylating PP1α87B/PP1 within its RVxF binding groove, POLO/PLK1 can decrease the affinity of PP1 for RVxF motifs, thereby influencing substrate dephosphorylation. The importance of this mechanism is demonstrated by its requirement for robust MPS1 activation and SAC signaling in *Drosophila* cells (Figure 7I). Though different organisms have evolved different circuits to control the SAC, the PLK1-PP1 regulatory axis may be conserved. It will be important in the future to verify whether this is its only critical role in mitosis and explore its implications in human disease.

## RESOURCE AVAILABILITY

### Lead contact

Further information and requests for resources and reagents should be directed to and will be fulfilled by the lead contact, Carlos Conde.

### Materials availability

All unique/stable reagents generated in this study will be made available on request with a completed Materials Transfer Agreement.

### Data and code availability

This study did not generate/analyze datasets/code.

## ACKNOWLEDGEMENTS

We thank Adrian Saurin (University of Dundee) and Reto Gassmann (i3S, University of Porto) for discussing the manuscript. We thank Geert Kops (Hubrecht Institute, The Netherlands) for the phospho-specific MPS1^T676h^ antibody (recognizes Drosophila MPS1^T490Ph^), Thomas Maresca (University of Massachusetts Amherst, USA) for the MAD1-EGFP construct, Helder Maiato (i3S/IBMC, Portugal) for the EGFP-CYCLIN B^WT^ and EGFP-CYCLIN B^NDG^ constructs and Gohta Goshima (Nagoya University, Japan) for the pAC-H2B-GFP and pAC-mCherry-α-Tubulin plasmids. This article is a result of the project NORTE-01-0145-FEDER-000051-Cancer Research on Therapy Resistance: From Basic Mechanisms to Novel Targets, supported by the Norte Portugal Regional Operational Programme (NORTE 2020), under the PORTUGAL 2020 Partnership Agreement, via the European Regional Development Fund (FEDER). MM was supported by the FCT PhD fellowship SFRH/BD/123306/2016. SC-S was supported by the FCT PhD fellowship SFRH/BD/136527/2018. IP is supported by the FCT PhD fellowship 2024.02612.BD. NL was supported by the FCT PhD fellowship SFRH/BD/136526/2018. JB and CC are supported by Scientific Employment Stimulus contracts (2023.07201.CEECIND and 2020.00067.CEECIND, respectively) from Fundação para a Ciência e a Tecnologia.

## AUTHOR CONTRIBUTIONS

Author contributions: M. Moura: Data curation, Formal analysis, Investigation, Methodology, Validation, Visualization, Writing - Original Draft, Writing - Review and Editing; J. Barbosa: Investigation, Validation, Visualization, Writing - Review and Editing; I. Pinto: Investigation, Validation, Visualization; N. Leça: Investigation; S. Cunha-Silva: Investigation, Writing - Review and Editing; P.D. Pedroso: Investigation, Software, Visualization; A.E. Verza: Investigation; S. Lemaire: Investigation; J. Duro: Investigation; M. Silva: Investigation; A. Oliveira: Investigation; R. Reis: Investigation; J. Nilsson: Investigation, Funding acquisition; C.E. Sunkel; Writing - Review and Editing, Funding acquisition; A. Musacchio: Writing - Review and Editing, Funding acquisition; M. Bollen: Investigation, Writing - Review and Editing, Funding acquisition; C. Conde: Conceptualization, Investigation, Data curation, Formal analysis, Funding acquisition, Project administration, Supervision, Writing - Original Draft, Writing - Review and Editing.

## CONFLICT OF INTEREST

The authors declare no competing financial interests.

## SUPPLEMENTAL INFORMATION

Figure S1 shows that PP1α87B inactivates MPS1 in metaphase regardless of CYCLIN A and CYCLIN B levels. Figure S2 shows that rendering POLO constitutively active increases PP1α87B T286 phosphorylation at metaphase kinetochores. Figure S3 shows that phosphorylation of PP1α87B at T286 modulates the activation status of MPS1 at prometaphase kinetochores. Figure S4 shows that kinetochore-tethering does not rescue the SAC-silencing delay associated with the PP1α87B^T286D^ mutation. Figure S5 shows that phosphorylation of T286/T288 does not affect the enzymatic activity of PP1α87B/PP1α/PP1γ, nor is it predicted to induce significant alterations of PP1α87B structure.

Video 1 and Video 2 show MAD1-mCherry kinetochore localization throughout mitosis in S2 cells expressing EGFP-CYCLIN B^WT^ or EGFP-CYCLIN B^NDG^, respectively. Video 3-7 show the mitotic progression of neuroblasts with the following genotypes: *w^1118^*(Video3), *pp1α87B^1^/pp1α87B^87Bg-3^* (Video 4), *pp1α87B^1^/pp1α87B^87Bg-3^;InscGal4>UAS-*PP1α87B^WT^-HA (Video 5), *pp1α87B^1^/pp1α87B^87Bg-3^;InscGal4>UAS-*PP1α87B^T286A^-HA (Video6) and *pp1α87B^1^/pp1α87B^87Bg-3^;InscGal4>UAS-*PP1α87B^T286D^-HA (Video 7)

**Document S1.** Figures S1 - S5 and Video legends

**Video S1.** The SAC is silenced even under elevated CDK1:CYCLIN B activity. Related to Figures 1C and 1D.

**Video S2.** The SAC is silenced even when CYCLIN B is rendered non-degradable. Related to Figure 1E and 1F.

**Video S3.** *w^1118^* neuroblasts faithfully segregate their chromosomes. Related to Figure 7C and 7D.

**Video S4.** *pp1α87B^1^/pp1α87B^87Bg-3^* neuroblasts mis-segregate their chromosomes. Related to Figure 7C and 7D.

**Video S5.** Neuroblasts expressing PP1α87B^WT^-HA in a *pp1α87B^1^/pp1α87B^87Bg-3^* genetic background faithfully segregate their chromosomes. Related to Figure 7C and 7D.

**Video S6.** Neuroblasts expressing PP1α87B^T286A^-HA in a *pp1α87B^1^/pp1α87B^87Bg-3^* genetic background exhibit lagging chromosomes in anaphase. Related to Figure 7C and 7D.

**Video S7.** Neuroblasts expressing PP1α87B^T286D^-HA in a *pp1α87B^1^/pp1α87B^87Bg-3^* genetic background exhibit asynchronous segregation of chromosomes in anaphase. Related to Figure 7C and 7D.

## MATERIAL AND METHODS

### S2 cell cultures, RNAi-mediated depletion, and drug treatments

The *Drosophila* S2-DGRC cell line (stock#006A9) was acquired from the Drosophila Genomics Resource Center, Indiana University, and was not independently authenticated. The cell lines routinely tested negative for mycoplasma contamination. *Drosophila* S2 cells were cultured at 25°C in Schneider’s Insect Medium (Sigma-Aldrich) supplemented with 10% fetal bovine serum (FBS). For RNAi-mediated depletion, 10^6^ S2 cells/mL on Schneider’s Insect Medium (Sigma-Aldrich) were plated on twelve-well plates and incubated with 30 µg of RNAi targeting MPS1 coding sequence (CDS), PP1α87B CDS, POLO CDS or PP1α87B 5’UTR and 3’UTR, for 120 h in 1.5 mL of Schneider’s insect medium (Sigma-Aldrich) supplemented with 10% FBS. The PCR products used as template to produce the indicated RNAis were amplified with following sets of primer pairs:

MPS1 CDS 5’TAATACGACTCACTATAGGGTCTTCCAAACACCTATGACCG3’/5’TAATACGACT CACTATAGGGCGTTTAGATATCCCTGCACCA3’;

PP1α87B 5’UTR 5’TAATACGACTCACTATAGGGAGACGGCAGTGTGGCAACATCAGCTA3’/5’TAATA CGACTCACTATAGGGAGAGTTTGCGTGCGAAAGTGTGGATCTG3’;

PP1α87B 3’UTR 5’AATACGACTCACTATAGGGAGATATCACACAACTGCAGCACCAC3’/5’TAATACG ACTCACTATAGGGAGAGTTTCATGCATCATTTGATGTGTTG3’;

PP1α87B CDS 5’TAATACGACTCACTATAGGGTCAGATCACATGAAGCTCATCC3’/5’TAATACGAC TCACTATAGGGCCTCCGAGAGCTGTACGTTT3’;

POLO 5’TAATACGACTCACTATAGGGTCAGATCACATGAAGCTCATCC3’/5’TAATACGAC TCACTATAGGGCTATAGCGCACTACGAACGC – 3’

RNAi was synthesized using the TranscriptAid T7 High Yield Transcription Kit (ThermoFisher) according to manufacture instructions. Cells were collected and processed at selected time points for immunofluorescence or time-lapse microscopy. When required, cells were subjected to several drug treatments before being collected and processed for the desired analysis. To promote microtubule depolymerization, cells were incubated with 30 µM colchicine (Sigma-Aldrich) for 2-16h. To inhibit POLO activity, 100 nM BI2536 (Boehringer Ingelheim) was added to cultured cells for 2h. When required, 20 µM MG132 (Calbiochem) were added to cultured cells to inhibit the proteasome.

### Constructs and S2 cell transfection

To generate the EGFP-PP1α87B construct for expression in S2 cells, PP1α87B cDNA was used as template to amplify the PP1α87B coding sequence by PCR using Phusion polymerase (ThermoFisher Scientific). The PCR product and the pMT-EGFP-C vector (Invitrogen, Carlsbad, CA) were digested with the restriction enzymes KpnI and XmaI (New England Biolabs) and the digested fragments were mixed in a proportion of 3:1 in the presence of T4 DNA Ligase (New England Biolabs) to obtain the PP1α87B coding sequence in frame with N-terminal EGFP under the regulation of a metallothionein promoter (pMT-EGFP-PP1α87B^WT^). The ligation product was used to transform DH5α competent bacteria, and cells were selected for the incorporation of plasmids. To clone the PP1α87B into the pENTR plasmid (Gateway Cloning System), PP1α87B cDNA was used as template to amplify the PP1α87B coding sequence by PCR using Phusion polymerase (ThermoFisher Scientific). The PCR product was inserted into the pENTR (Invitrogen, Carlsbad, CA) by FastCloning (Li et al., 2011). To generate pMT-EGFP-PP1α87B^T286A^, pMT-EGFP-PP1α87B^T286D^, pENTR-PP1α87B^T286A^, and pENTR-PP1α87B^T286D^, the recombinant pMT-EGFP-PP1α87B^WT^ and pENTR-PP1α87B^WT^ plasmids were used as templates to convert the codon corresponding to T286 into codons for alanine (A) and aspartate (D) by site-directed mutagenesis with primers harboring the desired mutations.

All the PCR amplifications were performed with Phusion polymerase (Thermo Fisher Scientific), and the resulting products were digested with the DpnI restriction enzyme (New England Biolabs) to remove the template DNA. The PCR products were used to transform DH5α competent bacteria, and cells were selected for the incorporation of plasmids. Recombinant plasmids pH-MIS12-GW-PP1α87B^WT^, pH-MIS12-GW-PP1α87B^T286A^, pH-MIS12-GW-PP1α87B^T286D^ were generated using the Gateway Cloning System (Invitrogen). The plasmids pHWG[blast]-POLO^WT^ and pHWG[blast]-POLO^T182D^ were previously characterized (Conde et al., 2013). To overexpress EGFP-MPS1 in S2 cells, the MPS1 coding sequence was cloned in frame with N-terminal EGFP under the regulation of a metallothionein promoter in the pMT-EGFP-C vector (Invitrogen, Carlsbad, CA) as previously described (Conde et al., 2013). The resulting pMT-EGFP-MPS1 was subsequently modified to generate pMT-PP1α87B^WT^-EGFP-MPS1. For that, the PP1α87B coding sequence was amplified by PCR and cloned into the pMT-EGFP-MPS1 plasmid by FastCloning (Li et al., 2011). The plasmids pMT-PP1α87B^T286A^-EGFP-MPS1, pMT-PP1α87B^T286D^-EGFP-MPS1 and pMT-PP1α87B^D62N^-EGFP-MPS1 were generated by site-directed mutagenesis with primers harboring the desired mutation. PCR reactions were performed with Phusion polymerase (Thermo Fisher Scientific) and pMT-PP1α87B^WT^-EGFP-MPS1 as template. PCR products were digested with DpnI restriction enzyme (New England Biolabs) and used to transform competent bacteria and selected for positives. The MAD1-EGFP construct under regulation of MAD1 native promoter cloned in the pMT backbone (Invitrogen) was a gift from Thomas Maresca (University of Massachusetts Amherst) and has been previously described (Cunha-Silva et al., 2020). The constructs EGFP-CYCLIN B^WT^ and EGFP-CYCLIN^NDG^ (Δ90 cyclinB) cloned in the pMT-EGFP-C vector (Invitrogen, Carlsbad, CA) were a gift from Helder Maiato and have been previously described (Afonso et al., 2014). The plasmids pAC-H2B-GFP and pAC-mCherry-α-Tubulin to drive constitutive expression of H2B-GFP and mCherry-α-Tubulin in S2 cells were a gift from Gohta Goshima (Nagoya University, Japan) and have been previously described (Goshima et al., 2007).

Stable transfection of the indicated vectors into S2 cells was performed using Effectene Transfection Reagent (Qiagen, Hilden, Germany), according to the manufacturer’s instructions. Stable cell lines were selected in medium with 25 µg/mL blasticidin. To induce the expression of EGFP-PP1α87B^WT^, EGFP-PP1α87B^T286A^, EGFP-PP1α87B^T286D^, EGFP-MPS1, PP1α87B^WT^-EGFP-MPS1, PP1α87B^T286A^-EGFP-MPS1, PP1α87B^T286D^-EGFP-MPS1, PP1α87B^D62N^-EGFP-MPS1, S2 cells were incubated with 100µM CuSO_4_ at 25°C for at least 8 hours before analysis. To induce the expression of POLO^WT^-EGFP, POLO^T182D^-EGFP, MIS12-EGFP-PP1α87B^WT^, MIS12-EGFP-PP1α87B^T286A^, MIS12-EGFP-PP1α87B^T286D^ and MIS12-EGFP, S2 cells were heat-shocked for 30 min at 37°C and allowed to rest for at least 8 h before being processed for immunofluorescence.

### Fly stocks

All fly stocks were obtained from Bloomington Stock Center unless stated otherwise. *w^1118^* was used as wild type control. PP1α87B mutant alleles, *pp1α87B^1^*and *pp1α87B^87Bg-3^*, have been described before (Axton et al., 1990; Kirchner et al., 2007). Fly stocks harboring WT, T286A or T286D versions of PP1α87B-HA were obtained by attP/attB PhiC31-mediated integration (BestGene Inc., USA). The pUASg-PP1α87B^WT^-HA, pUASg-PP1α87B^T286A^-HA and pUASg-PP1α87B^T286D^-HA constructs were generated using the Gateway Cloning System (Invitrogen).

### Rescue of viability experiments

UAS-PP1α87B-HA insertions on the third chromosome were recombined with PP1α87B^1^. These males were crossed to *arm*-GAL4;PP1α87B^Bg-3^ females to test for complementation of *pp1α87B^B1^/pp1α87B^87Bg-3^* mutant background. As a control for the quantification of the percentage of flies hatched, a similar cross scheme was performed for *arm-*GAL4-driven expression of UAS-lacZ in a wild type background.

### Live imaging of neuroblasts and S2 cells

For time-lapse imaging of *Drosophila* neuroblasts, brains from 3rd-instar larvae with indicated genotypes were dissected in PBS1× and placed on a drop of PBS1× in a coverslip. The preparation was gently squashed using a smaller coverslip, and excess media was removed. The sample was sealed with Halocarbon Oil 700 (Sigma-Aldrich). Images were obtained at 20-s intervals unless stated otherwise. For live imaging of S2 cells, cells were plated onto dishes with coverslip bottoms (MatTek Corporation, Ashland, MA, USA) previously treated with 0.2 mg/ml Concanavalin A (Sigma-Aldrich). All time-lapse images were acquired at 25°C with a spinning disc confocal system (Revolution; Andor) equipped with an electron-multiplying charge-coupled device camera (iXonEM+; Andor) and a CSU-22 unit (Yokogawa) based on an inverted microscope (IX81; Olympus). The microscope is served with two laser lines, 488 and 561 nm, for the excitation of EGFP and mCherry/mRFP, respectively. The system was driven by iQ software (Andor). Time-lapse imaging of z stacks with 0.8 µm steps covering the entire cell volume was collected, and image sequence analysis and video assembly were done with Fiji software (https://fiji.sc/). For quantification of MAD1-mCherry levels at kinetochores throughout mitosis, the mean fluorescence intensity of MAD1-mCherry was measured within specific predefined regions of interest (ROI) corresponding to the areas of individual kinetochores, which were identified based on the accumulation of EGFP-CYCLIN B^WT^ or EGFP-CYCLIN B^NDG^. The mean fluorescence intensity of cytoplasmic MAD1-mCherry was also determined. After subtraction of extracellular background signal to kinetochore and cytoplasm signals, the ratio (kinetochore–background)/(cytoplasm–background) was determined and used as a measure of MAD1-mCherry accumulation at kinetochores. The mean MAD1-mCherry ratio per cell was calculated for all frames and plotted against time. The integrated intensities of EGFP-CYCLIN B^WT^ or EGFP-CYCLIN B^NDG^ fluorescence were measured in the same frames and plotted as normalized signals relative to the averaged signal of early prometaphase frames.

### Immunofluorescence analysis of neuroblasts and S2 cells

For immunofluorescence analysis of *Drosophila* neuroblasts, third-instar larval brains were dissected in PBS and incubated with 50 µM colchicine for 90 min. Afterward, the brains were fixed in 1.8% formaldehyde (Sigma-Aldrich) and 45% glacial acetic acid for 5 min, squashed between slide and coverslip, and immersed in liquid nitrogen. Subsequently, coverslips were removed, and the slides were incubated in cold ethanol for 10 min and washed in PBS with 0.1% Triton X-100. Immunostaining of neuroblasts was performed as previously described (Moura et al., 2017). For scoring aneuploidy levels, larval brains were subjected to a 10 min hypotonic shock before undergoing fixation.

For immunofluorescence analysis of S2 cells, 105 cells were centrifuged onto slides for 5 min at 1,500 rpm (Cytospin 2, Shandon). Afterward, cells were fixed in 4% PFA in PBS for 12 min and then extracted for 8 min with 0.1% Triton X-100 in PBS. Alternatively, cells were simultaneously fixed and extracted in 3.7% formaldehyde (Sigma-Aldrich) and 0.5% Triton X-100 in PBS for 10 min followed by three washing steps of 5 min with PBST (PBS with 0.05% Tween20). Immunostaining of S2 cells was performed as previously described (Moura et al., 2017). Images were collected on an inverted Laser Scanning Confocal Microscope Leica TCS SP8 (Leica Microsystems, Germany) with the PL APO 63x/1.30 Glycerol objective, using the LAS X software (Leica Microsystems, Germany). All images were analyzed using the Fiji software (https://fiji.sc/). For immunofluorescence quantification, the mean pixel intensity was obtained from maximum projected raw images acquired with fixed exposure acquisition settings. The mean fluorescence intensity was quantified for individual kinetochores, selected manually based on CID or SPC105R staining. The area of the region of interest (ROI) was predefined so that each single kinetochore could fit into it. After subtraction of background intensities, estimated from extracellular areas, the intensity was determined relative to CID, SPC105R or EGFP references and averaged over multiple kinetochores.

### Production and purification of recombinant PP1 for enzymatic assays

To generate the MBP-PP1α87B construct for expression in bacteria, PCR products harbouring PP1α87B coding sequence were cloned into StuI/XbaI sites of pMal-c2 (New England Biolabs) vector. To generate the MBP-PP1α87B^T286D^ construct, the recombinant pMal-c2-PP1α87B^WT^ plasmid was used as template to convert the codon corresponding to T286 into codons for aspartate (D) by site-directed mutagenesis. The recombinant constructs were used to transform BL21-star competent cells and protein expression induced with 0.1 mM isopropyl β-D-1-thiogalactopyranoside (IPTG) at 15°C, overnight. Lysates were sonicated and clarified by centrifugation at 4 °C. Recombinant MBP-PP1α87B^WT^ or MBP-PP1α87B^T286D^ were purified with amylose magnetic beads (New England Biolabs) and eluted with 10 mM Maltose, 200 mM NaCl, 20 mM Tris-HCl pH 7.4, 1 mM EDTA and 1 mM DTT.

Constructs for human GST-PP1α were cloned into pGEX-2TK vector and expressed in BL21 gold cells at 37 °C in LB medium with 100 µg/mL ampicillin. The cells were grown until an OD600 of ≈ 0.6, then protein expression was induced for 20 h at 18 °C with 1 mM IPTG. Cells were lysed in 10 volumes of a buffer containing 50 mM Tris at pH 7.4, 0.3 M NaCl, 1 mM DTT, 0.5% Triton X-100, 1 mM MnCl2, 0.05 mg/mL lysozyme and protease inhibitors (1 mM PMSF, 1 mM benzamidine, 5 ug/mL leupeptin). Lysates were cleared by centrifugation (20 min at 15,000 × g) and the supernatant was loaded for 1 h at 4 °C onto glutathione agarose. The loaded glutathione-agarose beads were washed with 50 mM Tris at pH 7.4, 150 mM NaCl, 1 mM DTT, 1 mM MnCl2, and eluted with 0.1 M Tris at pH 8.0, 1 mM DTT, 1 mM MnCl2 and 10 mM reduced glutathione. The eluted fractions were pooled and dialyzed overnight at 4°C against 50 mM Tris at pH 7.5, 0.1 M NaCl, 1 mM DTT, 1mM MnCl2 and 60% glycerol, and stored at -80 °C.

His-tagged PP1γ was expressed in BL21(DE3) overnight at 18 degrees. The cell pellet was resuspended in lysis buffer (50 mM NaP pH 7,5, 300 mM NaCl, 10 mM imidazole, 10% glycerol 0,5 mM TCEP, benzonase, protease inhibitor cocktail EDTA free) and lysed by sonication. Following clarification by centrifugation the cell extract was loaded on a 5 mL HiTrap Ni column and then washed with lysis buffer followed by a wash with lysis buffer containing 30 mM imidazole. Elution was achieved using an imidazole gradient from 30-500 mM and peak fractions collected and pooled. The sample was dialysed into TEV cleavage buffer (50 mM NaP pH 7.5, 300 mM NaCl, 10% glycerol, 0,5 mM TCEP) and cleaved overnight with TEV. The sample was run on a 5 ml HiTrap Ni column and run through collected and concentrated. The sample was run on a Superdex 200 16/60 column equilibrated with 50 mM NaP pH=7,5, 150 mM NaCl, 10% glycerol, 0,5 mM TCEP and peak fractions pooled.

### *In vitro* phosphatase assays

*In vitro* phosphatase activity assays with the fluorogenic 6,8-difluoro-4-methylumbelliferyl phosphate (DiFMUP) substrate were conducted at 30 °C in 96-well plates in a final volume of 100 µl and at a final substrate concentration of 200 µM. The assay buffer contained 50 mM HEPES, pH 7.5, 100mM NaCl, 0.01% NP-40, 100 μM MnCl_2_ and 2 mM DTT. Phosphatase activities were measured at 30 °C within a timeframe ranging from 36 cycles of 5 min each on a Synergy 2 reader at an excitation of 360 nm and emission of 460 nm.

### *In vitro* kinase assays

For *in vitro* kinase assays, recombinant 6xHis-PP1γ^WT^ or 6x His-PP1γ^T288A^ at a final concentration of 450 nM were incubated with increasing concentrations (50 nM, 100 nM, 200 nM) of 6xHis-PLK1 (SignalChem) for 30 min. Reactions were carried out at 30°C in the presence of Phosphatase Inhibitors (Cocktail 3, Sigma-Aldrich) and 100 μM of Okadaic Acid in a total volume of 30 mL kinase reaction buffer (5 mM MOPS, pH 7.2, 2.5 mM Δ-glycerol-phosphate, 5 mM MgCl2, 1 mM EGTA, 0.4 mM EDTA, 0.05 mM DTT and 100 μM ATP). Reactions were stopped by addition of Laemmli sample buffer (4% SDS, 10% Δ-mercaptoethanol, 0.125 M Tris-HCl, 20% glycerol, 0.004% bromophenol blue) and heated for 5 min at 95°C. Peptides were resolved in 8% SDS-PAGE. The phosphorylation of PP1γ at T288 was detected by western blotting with a phospho-specific antibody.

### Production and purification of recombinant proteins for pull-down assays

To generate 6xHis-PP1γ^WT^ and 6xHis-PP1γ^T288D^ constructs for expression in bacteria, the PP1γ^WT^ and PP1γ^T288D^ coding sequences were cloned into the HindIII/XhoI sites of pET30a (+) vector (Novagen, Darmstadt, Germany). The PP1γ^T288D^ coding sequence was obtained by converting the T288 codon of PP1γ^WT^ into aspartate (D) via site-directed mutagenesis. The recombinant plasmids, pET30a-PP1γ^WT^ and pET30a-PP1γ^T288D^, were amplified by PCR to remove a fragment containing the thrombin site and S-tag. The final pET30a-PP1γ^WT^ and pET30a-PP1γ^T288D^ plasmids were then used as templates to introduce mutations that render the PP1γ protein catalytic-dead. To do so, the codon for D64 was converted into a codon for asparagine (N) via site-directed mutagenesis, yielding pET30a-PP1γ^D64N^ and pET30a-PP1γ^D64N/T288D^. To generate the MBP-MPS1^104-330^ construct for expression in bacteria, a PCR product containing the sequence encoding amino acids 104-330 of MPS1 from Drosophila melanogaster was inserted into the pMal-c2 vector (New England Biolabs) by FastCloning (Li et al., 2011). To generate the MBP-MPS1^104-330/4A^ construct, the recombinant pMal-c-MPS1^104-330^ was used as a template to convert the codons corresponding to K231, V232, L233, and F234 into codons for alanine (A), by site-directed mutagenesis with primers harboring the desired mutations. All the PCR amplifications were performed with Phusion polymerase (Thermo Fisher Scientific), and the resulting products were digested with the DpnI restriction enzyme (New England Biolabs) to remove the template DNA. The ligation and PCR products were used to transform DH5α competent bacteria, and cells were selected for the incorporation of plasmids. Selected recombinant constructs were used to transform BL21-star competent cells for protein expression.

All protein purification steps were carried out on ice or at 4°C. Cells expressing 6xHis-PP1γ^D64N^ or 6xHis-PP1γ^D64N/T288D^ were resuspended in purification buffer (50 mM Hepes pH 7.5, 300 mM NaCl, 5% glycerol, 1 mM TCEP), with the addition of DNAse (Roche, Basel, Switzerland) and Protease inhibitors (Serva Electrophoresis GmbH, Heidelberg, Germany), and sonicated to induce cell rupture. The total lysate was clarified by centrifugation at 85,000×g for 30 minutes and then mixed with 2 mL cOmplete His-tag NiNTA beads (Roche) pre-equilibrated in purification buffer. After an incubation under agitation of 2 hours, beads were washed on a 10-mL gravity column (Thermo Fisher Scientific, Waltham, US-MA) with 40 mL of purification buffer and 50 mL of wash buffer (50 mM Hepes pH 7.5, 300 mM NaCl, 5% glycerol, 1 mM TCEP, 20 mM imidazole). Protein was eluted in three 2-mL fractions with 300 mM Imidazole in purification buffer. The first fraction was discarded while the other two were cleared by centrifugation at 16,900×g, and the supernatant was directly injected in a Superdex 200 10/600 (Cytiva, Marlborough, US-MA). The eluted fractions were inspected by SDS-PAGE and Coomassie blue staining, and the purest and most concentrated fractions were collected, spun down at 16900 g and directly snap-frozen and stored at -80°C. As further protein concentration caused precipitation, it was avoided to increase the final yield of this specific protein.

MBP-MPS1^104-330/WT^ and MPS1^104-330/4A^ were obtained by over-expression in 1 L culture of BL21CodonPlus(DE3)-RIL cells (Agilent Technologies, Santa Clara, California, United States) grown in Terrific Broth (TB). Expression was induced with IPTG 0.1 mM and cells were grown for 18 hours at 18°C. Cells were pelleted, snap-frozen and stored at -80°C until purification. All purification steps were performed at 4°C or on ice. A resuspension of cells and purification buffer (20 mM Tris pH 8, 500 mM NaCl, 1 mM TCEP) was sonicated to lyse cells and centrifuged at 85,000×g for 30 minutes. The resulting clarified lysate was incubated with 4 mL slurry Amylose beads (New England Biolabs, Ipswich, US-MA) and kept under constant agitation for 2 hours. Beads were washed with 30 mL purification buffer and further incubated under constant agitation for 15 hours in 15 mL purification buffer. Beads were then washed with additional 30 mL buffer and protein was eluted by addition of maltose powder (Carl Roth GmbH, Karlsruhe, Germany) to purification buffer at a 10 mM final concentration. Protein was concentrated and injected in a Superdex 200 16/600 (Cytiva). Fractions were inspected for purity via SDS-PAGE gel and Coomassie Brilliant Blue staining. The purest fractions were pooled together and concentrated, aliquoted, snap-frozen and stored at -80°C.

### Pull-down assays

Amylose-resin pull-down assays were performed to evaluate the interaction between MBP-MPS1^104-330/WT^ or MPS1^104-330/4A^ and 6xHis-PP1γ^WT^ or 6xHis-PP1γ^T288D^. The purified recombinant proteins were diluted in reaction buffer consisting of 150 mM NaCl, 50 mM HEPES pH 7.5, 1 mM TCEP, 5% Glycerol and 0.03% Tween20, and mixed at a final concentration of 5 µM for MBP-MPS1 constructs and 2 µM for 6xHis-PP1 constructs, in a total volume of 40 µL. The amylose beads slurry (New England BioLabs) was equilibrated in buffer and 20 µL were added to each mix for a 4°C incubation of 1 h. The input consists of 10 µL sample-beads mix with 40 µL of 2× SDS sample loading buffer. The samples were washed four times with 500 µL of the same buffer and centrifuged at 1 000 g for 1 min at 4°C to separate beads from the washing solution. After the last washing step, 50 µL of 2× SDS sample loading buffer was added to the dry beads to obtain the bound fraction.

### Immunoblotting and gel staining

Coomassie staining of proteins resolved by SDS–PAGE was first conducted to determine comparable amounts of recombinant protein in solution to be used in kinase assays and pull-down experiments. For immunoblotting analysis, resolved proteins were transferred to a nitrocellulose membrane, using the iBlot Dry Blotting System (Thermo Fisher Scientific) according to the manufacturer’s instructions. Membranes were incubated at room temperature for at least 1 h in blocking solution (5% powder milk in PBS1×, 0.05% Tween 20). All primary and secondary antibodies were diluted in the blocking solution. Membranes were incubated with primary antibody solutions overnight at 4°C under constant stirring and washed three times in PBS1×, 0.05% Tween 20 for 10 min each. Then, membranes were incubated with secondary antibody solutions for 1 h at room temperature under constant stirring. Secondary antibodies were conjugated to horseradish peroxidase (Jackson Immuno Research, UK). The blots were visualized by ECL detection and exposure to X-ray films (Fuji Medical X-Ray Film). For immunoblotting analysis of the pull-down assays, resolved proteins were transferred to a nitrocellulose membrane, using the TransBlot Turbo transfer system (BioRad Laboratories, Hercules, US-CA) according to the manufacturer’s instructions. Membranes were incubated for at least 1 h at room temperature in blocking solution (5% powder milk in TBS 1×, 0.1% Tween 20). Primary antibodies were diluted in the blocking solution. Membranes were incubated with primary antibody solutions overnight at 4°C under constant stirring and washed three times in TBS1×, 0.1% Tween 20 for 10 min each. Then, membranes were incubated with secondary antibody dissolved in TBS1×, 0.1% Tween 20 for 1 h at room temperature under constant stirring. Secondary antibodies were conjugated to horseradish peroxidase (Amersham-Cytiva-catalog number NXA931-1ML) and used at 1:10000 dilution. The blots were visualized by ECL detection and images were acquired with the ChemiDoc MP System (Bio-Rad) using Image Lab 5.1 software.

### Antibodies

Rabbit polyclonal anti-PP1-phospho(ph)Thr288 was raised against a phosphorylated peptide MMSVDE-pT-LMCSFQ, generated by Eurogentec, Seraing, Belgium. The following primary antibodies were used for immunofluorescence studies: rat anti-CID (Rat4) used at 1:250, chicken anti-GFP (ab13970; Abcam) used at 1:2000, mouse anti-HA (26183, Invitrogen) used at 1:500, rabbit anti-phosphorylated Thr716-MAD1 (MAD1^T716Ph^; Allan et al., 2020) used at 1:1000, rabbit anti-phosphorylated Thr676-MPS1 (MPS1^T676Ph^; a gift from Geert Kops, Hubrecht Institute, Utrecht, Netherlands; Jelluma et al., 2008) used at 1:20000, rabbit anti-phosphorylated Thr288-PP1 used at 1:500, rat anti-Spc105R used at 1:250. The following secondary antibodies were used for immunofluorescence studies: Alexa anti-chicken 488 used at 1:2000, Alexa anti-mouse 488 used at 1:2000, Alexa anti-mouse 568 used at 1:2000, Alexa anti-rabbit 488 used at 1:2000, Alexa anti-rabbit 568 used at 1:2000 and Alexa anti-rat 647 used at 1:1000. The following primary antibodies were used for Western blotting: mouse anti-His-Tag (05-949, Milipore) used at 1:2500, rabbit anti-phosphorylated Thr288-PP1 used at 1:250 and mouse anti-PP1γ (Santa Cruz, #sc-515943, Clone E-4), used at 1:1000. HRP-conjugated anti-mouse and anti-rabbit secondary antibodies (Jackson ImmunoResearch) were used according to the manufacturer’s instructions.

### AlphaFold modeling

The ColabFold v1.5.5 of AlphaFold 2 (Jumper et al., 2021) was used to model the structure of *Drosophila* PP1α87B (P12982) and the impact of mutating T286 to alanine (T286A) or aspartate (T286D) on the overall structure. The structure of wild type PP1α87B (WT), T286A and T286D mutants was predicted with the following parameters: use_templates: false, num_relax: 1, relax_max_iterations: 200, relax_tolerance: 2.39, relax_stiffness: 10.0, relax_max_outer_iterations: 3, msa_mode: mmseqs2_uniref_env, model_type: alphafold2_ptm, num_models: 5, num_recycles: 6, recycle_early_stop_tolerance: null, num_ensemble: 1, rank_by: plddt, max_seq: 512, max_extra_seq: 5120, pair_mode: unpaired_paired, pairing_strategy: greedy.

AlphaFold 3 (Abramson et al, 2024) was used to model the structure of PP1α87B and the impact of T286 phosphorylation (T286Ph) on the interaction with the RVxF motif of *Drosophila* MPS1. The PP1α87B structure was predicted by inputting the whole PP1α87B sequence as a single copy ‘Protein’ with the Fe^2+^ and the Zn^2+^ ions as single copy ‘Ion’ in the catalytic pocket. The binding models were also obtained with the whole PP1α87B sequence, where in this case either as its wild type unmodified version or with a phospho-threonine PTM added onto the T286 residue (T286Ph). At the same time, a single copy of *Drosophila* MPS1 ^226^DFRAQKVLFQTPMT^239^, was added to predict its binding to both PP1α87B variants. Clash Score analysis was assessed in SwissModel for each of the five output models to determine quality parameters for each condition. Those can be found in Figure S5. All figures of the predicted structures, either for AlphaFold2 or AlphaFold3, were further processed using the Mol* 3D Viewer from RCSB PDB, where colouring and representations were added.

### Statistical analysis

All statistical analysis was performed with GraphPad Prism V10.0 (GraphPad Software, Inc.). Values were considered statistically different whenever P < 0.05. P values: ns, not significant; *< 0.05; **< 0.01; ***< 0.001; ****< 0.0001. The normality of the samples was determined with a D’Agostino & Pearson test. Statistical analysis for two sample comparison, with normal or non-normal distribution, was performed with a t-test or Mann-Whitney test, respectively. For multiple group comparison a parametric one-way analysis of variance (ANOVA) or a nonparametric ANOVA (Kruskal-Wallis) was used for samples with normal or non-normal distribution, respectively. All pairwise multiple comparisons were subsequently analyzed using either Student t-test (parametric) or Dunn’s (nonparametric) tests.

## Notes

### Competing Interest Statement

The authors have declared no competing interest.

