## Supplemental Information for "POLO kinase inhibits Protein Phosphatase 1 to promote the Spindle Assembly Checkpoint and prevent aneuploidy"

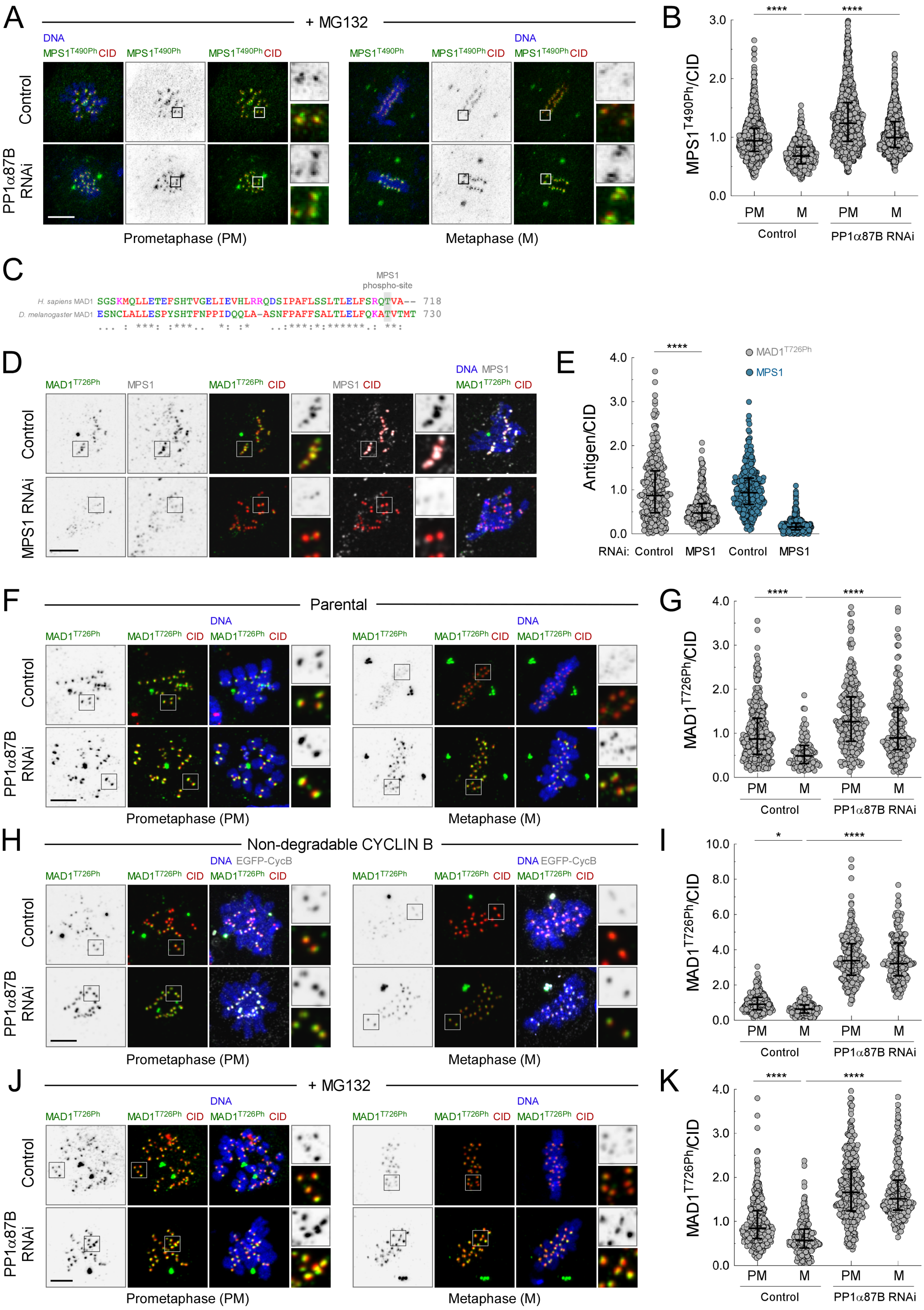


Figure S1 Moura et al

**Figure S1. PP1α87B-mediated inactivation of MPS1 in metaphase is impervious to CDK1 activity. Related to Figure 1.** (A,B) Representative immunofluorescence images (A) and corresponding quantifications (B) of MPS1 T490 (MPS1$^{T490Ph}$) phosphorylation levels at kinetochores of control and PP1α87B-depleted S2 cells treated with MG132 for 2 hours ($n \geq 593$ cells for each condition). MPS1$^{T490Ph}$ fluorescence intensity was determined relative to the CID signal. (C) Clustal W alignment of *H. sapiens* and *Drosophila* MAD1 orthologues. The conserved C-terminus threonine phosphorylated by MPS1 is highlighted in grey shading. (D,E) Representative immunofluorescence images (D) and corresponding quantifications (E) of MPS1 and MAD1 T726 phosphorylation (MAD1$^{T726Ph}$) levels at unattached kinetochores of control and MPS1-depleted S2 cells incubated with MG132 (1 hour) and colchicine (3 hours). MAD1$^{T726Ph}$ and MPS1 fluorescence intensities were determined relative to CID signal ($n \geq 326$ kinetochores for each condition). (F-K) Representative immunofluorescence images (F,H,J) and corresponding quantifications (G,I,K) of MAD1 T726 phosphorylation (MAD1$^{T726Ph}$) levels at kinetochores of control and PP1α87B-depleted S2 cells in prometaphase and metaphase, Asynchronous parental S2 cells (F,G), cells expressing EGFP-CYCLIN B$^{NDG}$ (H,I) and MG132-treated (2 hours) parental S2 cells were analyzed for MAD$^{T726Ph}$ fluorescence intensity relative to CID signal ($n \geq 109$ cells for each condition). Data information: data in B,E,G,I,K are presented as median plus interquartile range; asterisks indicate that differences between mean ranks are statistically significant, $*p<0.05$, $****p<0.0001$ ns, non-significant (Kruskal-Wallis, Dunn's multiple comparison test in B, G, I and K and Mann-Whitney U test in E). Scale bars: 5μm.

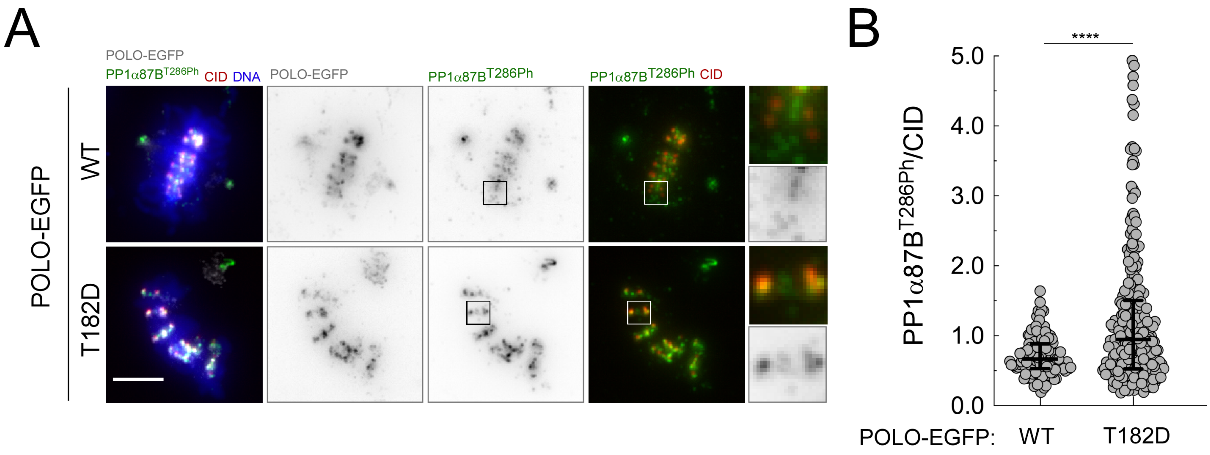


Figure S2 Moura et al

**Figure S2. Mimicking constitutive activation of POLO increases PP1α87B T286 phosphorylation at metaphase kinetochores. Related to Figure 2.** (A,B) Representative immunofluorescence images (A) and corresponding quantifications (B) of PP1α87B T286 phosphorylation (PP1α87B$^{T286Ph}$) levels at metaphase kinetochores of S2 cells expressing POLO$^{WT}$-EGFP or Polo$^{T182D}$-EGFP. PP1α87B$^{T286Ph}$ fluorescence intensity was determined relative to CID signal ($n \geq 134$ kinetochores for each condition). Data information: data in B is shown as median plus interquartile range; asterisks indicate that differences between mean ranks are statistically significant, ****$p<0.0001$ (Mann-Whitney U test). Scale bars: 5µm.

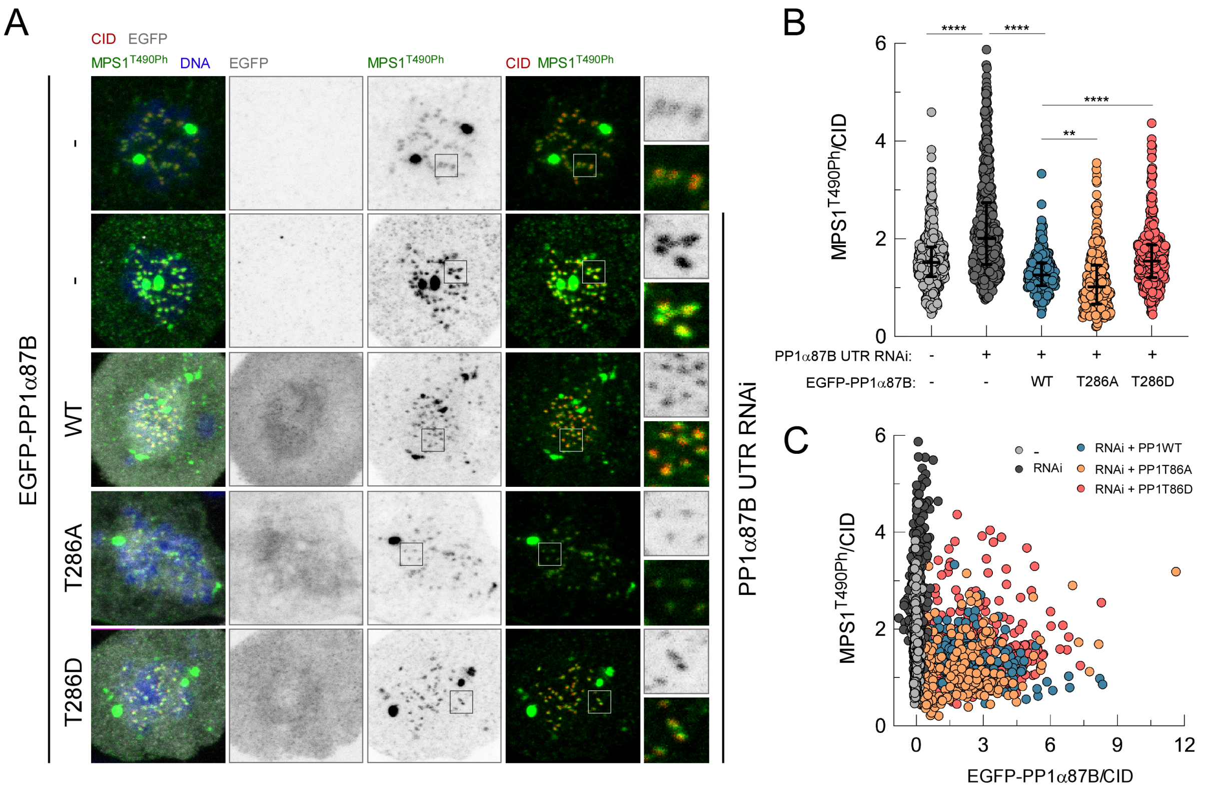


Figure S3 Moura et al

**Figure S3. Phosphorylation of PP1α87B at T286 controls MPS1 activation at prometaphase kinetochores. Related to Figure 4.** (A-C) Representative immunofluorescence images (A) and corresponding quantifications (B) of MPS1$^{T490Ph}$ levels at the kinetochores of control and PP1α87B-depleted S2 cells in prometaphase expressing the indicated EGFP-PP1α87B transgenes. (C) MPS1$^{T490Ph}$ levels in B plotted over EGFP-PP1α87B levels at kinetochores. MPS1$^{T490h}$ and EGFP fluorescence intensities were determined relative to CID signal ($n \geq 300$ kinetochores for each condition). Data information: data in B are presented as median with interquartile range; asterisks indicate that differences between mean ranks are statistically significant, **$p<0.01$, ****$p<0.0001$ (Kruskal-Wallis, Dunn's multiple comparison test). Scale bars: 5μm.

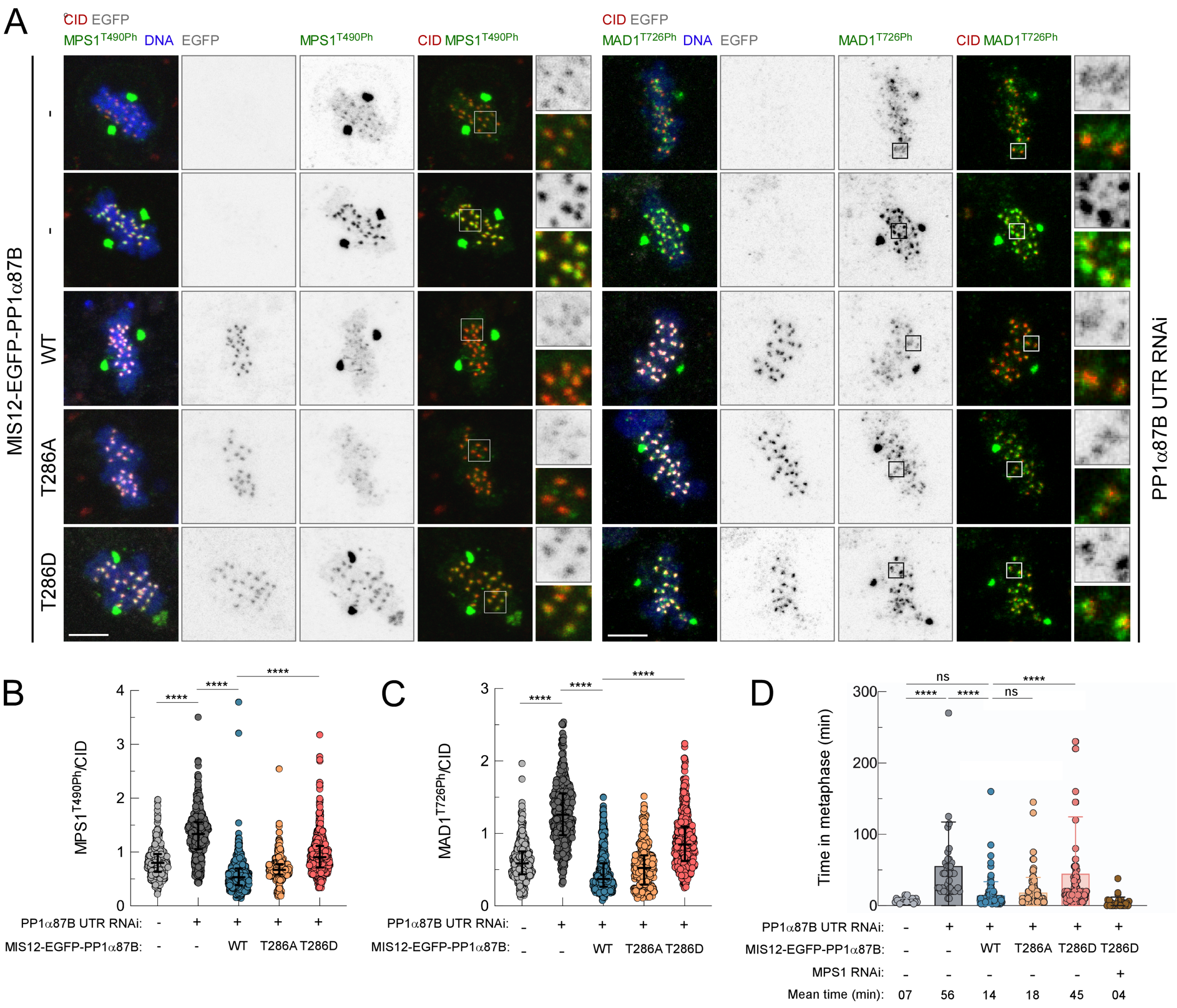


Figure S4 Moura et al

**Figure S4. *Drosophila* S2 cells expressing kinetochore-tethered phosphomimetic PP1α87B$^{T286D}$ are delayed in SAC silencing. Related to Figure 5.** (A-C) Representative immunofluorescence images (A) and corresponding quantifications (B,C) of MPS1 T490 phosphorylation (MPS1$^{T490Ph}$) or MAD1 T726 phosphorylation (MAD1$^{T726Ph}$) levels at kinetochores of control and PP1α87B-depleted S2 cells in metaphase expressing the indicated MIS12-EGFP-PP1α87B transgenes. MPS1$^{T490h}$ and MAD1$^{T726h}$ fluorescence intensities were determined relative to CID signal ($n \geq 295$ kinetochores for each condition in B and $n \geq 373$ kinetochores for each condition in C). (D) Metaphase duration of control and PP1α87B-depleted S2 cells in metaphase expressing the indicated MIS12-EGFP-PP1α87B transgenes. The time in metaphase was monitored by time-lapse microscopy and corresponds to the length of time measured between the frame when all chromosomes align at the metaphase plate and the frame when anaphase onset occurs ($n \geq 23$ cells for each condition). Data information: data in B and C are presented as median with interquartile range and data in D are presented as mean with standard deviation; asterisks indicate that differences between mean ranks are statistically significant, ****$p<0.0001$; ns, non-significant (Kruskal-Wallis, Dunn's multiple comparison test). Scale bars: 5μm.

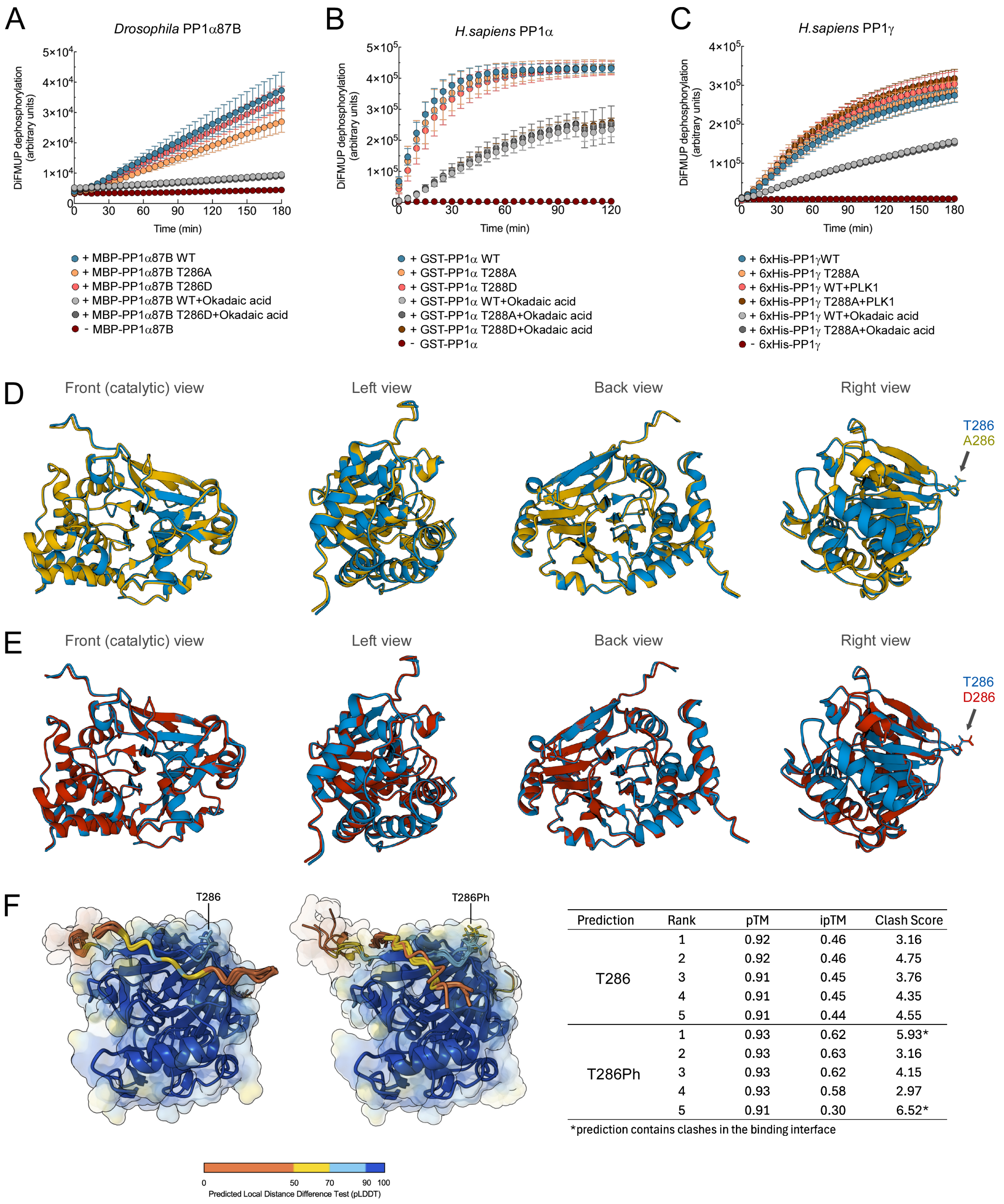


| Prediction | Rank | pTM | ipTM | Clash Score |
|---|---|---|---|---|
| T286 | 1 | 0.92 | 0.46 | 3.16 |
| | 2 | 0.92 | 0.46 | 4.75 |
| | 3 | 0.91 | 0.45 | 3.76 |
| | 4 | 0.91 | 0.45 | 4.35 |
| | 5 | 0.91 | 0.44 | 4.55 |
| T286Ph | 1 | 0.93 | 0.62 | 5.93* |
| | 2 | 0.93 | 0.63 | 3.16 |
| | 3 | 0.93 | 0.62 | 4.15 |
| | 4 | 0.93 | 0.58 | 2.97 |
| | 5 | 0.91 | 0.30 | 6.52* |

*prediction contains clashes in the binding interface

Figure S5 Moura et al

**Figure S5. Phosphorylation of T286/T288 does not affect the catalytic activity of PP1α87B/PP1 but decreases its ability to bind to the RVxF motif of MPS1. Related to Figure 6.** (A-C) EnzChek phosphatase assays with purified *Drosophila* PP1α87B (A), *H. sapiens* PP1α (B) and *H. sapiens* PP1γ (C). DiFMUP dephosphorylation by wild type (WT), unphosphorylatable (T286A/T288A) and phosphomimetic (T286D/T288D) versions of recombinant MBP-PP1α87B, GST- PP1α and 6xHis-PP1γ was monitored over time in the indicated conditions. All recombinant proteins were produced in and purified from bacteria. (D,E) Superposed AlphaFold 2 models of *Drosophila* PP1α87B WT (blue) with PP1α87B T286A (yellow) and PP1α87B T286D (red) (E) showing that the T286A and T286D mutations are not predicted to induce major alterations in the structural backbone of PP1α87B.

(F) Overlap of five AlphaFold 3 models with pLDDT colouring of the interaction between the RVxF motif ($^{231}$KVLF$^{234}$) of *Drosophila* MPS1 and PP1α87B or PP1α87B phosphorylated on T286 (T286Ph) and their respective predicted template modelling (pTM), interface predicted template modelling (ipTM) and Clash Scores values.

### VIDEO LEGENDS

**Video S1. The SAC is silenced even under elevated CDK1:CYCLIN B activity. Related to Figures 1C and 1D.** Mitotic progression of *Drosophila* S2 cells expressing MAD1-mCherry and EGFP-CYCLIN B$^{WT}$. Merge colors of MAD1-mCherry (red) and EGFP-CYCLIN B$^{WT}$ (green) channels are presented at the left. The MAD1-mCherry channel is displayed at the center and the EGFP-CYCLIN B$^{WT}$ channel is at the right. Frames were acquired every 30 sec. Time 0 corresponds to anaphase onset.

**Video S2. The SAC is silenced even when CYCLIN B is rendered non-degradable. Related to Figure 1E and 1F.** Mitotic progression of *Drosophila* S2 cells expressing MAD1-mCherry and EGFP-CYCLIN B$^{NDG}$. Merge colors of MAD1-mCherry (red) and EGFP-CYCLIN B$^{NDG}$ (green) channels are presented at the left. The MAD1-mCherry channel is displayed at the center and the EGFP-CYCLIN B$^{NDG}$ channel is at the right. Frames were acquired every 30 sec. Time 0 corresponds to the first frame where maximal inter-kinetochore distance was achieved, as determined based on EGFP-Cyclin B$^{NDG}$ signal for kinetochore reference.

**Video S3. *w*$^{1118}$ neuroblasts faithfully segregate their chromosomes. Related to Figure 7C and 7D.** Mitotic progression of *Drosophila* *w*$^{1118}$ neuroblasts expressing H2Av-mRFP and EGFP-α-TUBULIN. Merge colors of H2Av-mRFP (red) and EGFP-α-TUBULIN (green) channels are shown. Frames were acquired every 20 sec. Time 0 corresponds to nuclear envelope breakdown.

**Video S4.** ***pp1α87B$^{1}$/pp1α87B$^{87Bg-3}$*** **neuroblasts mis-segregate their chromosomes. Related to Figure 7C and 7D.** Mitotic progression of *Drosophila pp1α87B$^{1}$/pp1α87B$^{87Bg-3}$* neuroblasts expressing H2Av-mRFP and EGFP-α-TUBULIN. Merge colors of H2Av-mRFP (red) and EGFP-α-TUBULIN (green) channels are shown. Frames were acquired every 20 sec. Time 0 corresponds to nuclear envelope breakdown.

**Video S5. Neuroblasts expressing PP1α87B$^{WT}$-HA in a** ***pp1α87B$^{1}$/pp1α87B$^{87Bg-3}$*** **genetic background faithfully segregate their chromosomes. Related to Figure 7C and 7D**. Mitotic progression of *Drosophila pp1α87B$^{1}$/pp1α87B$^{87Bg-3}$* neuroblasts expressing PP1α87B$^{WT}$-HA, H2Av-mRFP and EGFP-α-TUBULIN. Merge colors of H2Av-mRFP (red) and EGFP-α-TUBULIN (green) channels are shown. Frames were acquired every 20 sec. Time 0 corresponds to nuclear envelope breakdown.

**Video S6. Neuroblasts expressing PP1α87B$^{T286A}$-HA in a** ***pp1α87B$^{1}$/pp1α87B$^{87Bg-3}$*** **genetic background exhibit lagging chromosomes in anaphase. Related to Figure 7C and 7D**. Mitotic progression of *Drosophila pp1α87B$^{1}$/pp1α87B$^{87Bg-3}$* neuroblasts expressing PP1α87B$^{T286A}$-HA, H2Av-mRFP and EGFP-α-TUBULIN. Merge colors of H2Av-mRFP (red) and EGFP-α-TUBULIN (green) channels are shown. Frames were acquired every 20 sec. Time 0 corresponds to nuclear envelope breakdown.

**Video S7. Neuroblasts expressing PP1α87B$^{T286D}$-HA in a** ***pp1α87B$^{1}$/pp1α87B$^{87Bg-3}$*** **genetic background exhibit asynchronous segregation of chromosomes in anaphase. Related to Figure 7C and 7D.** Mitotic progression of *Drosophila pp1α87B$^{1}$/pp1α87B$^{87Bg-3}$* neuroblasts expressing PP1α87B$^{T286D}$-HA, H2Av-mRFP and EGFP-α-TUBULIN. Merge colors of H2Av-mRFP (red) and EGFP-α-TUBULIN (green) channels are shown. Frames were acquired every 20 sec. Time 0 corresponds to nuclear envelope breakdown.
